# HiDDEN: A machine learning label refinement method for detection of disease-relevant populations in case-control single-cell transcriptomics

**DOI:** 10.1101/2023.01.06.523013

**Authors:** Aleksandrina Goeva, Michael-John Dolan, Judy Luu, Eric Garcia, Rebecca Boiarsky, Rajat M Gupta, Evan Macosko

## Abstract

In case-control single-cell RNA-seq studies, sample-level labels are transferred onto individual cells, labeling all case cells as affected, but only a small fraction of them may actually be perturbed. Here, using simulations, we demonstrate that the standard approach to single cell analysis fails to isolate the subset of affected case cells and their markers when either the affected subset is small, or when the strength of the perturbation is mild. To address this fundamental limitation, we introduce HiDDEN, a computational method that refines the case-control labels to accurately reflect the perturbation status of each cell. We show HiDDEN’s superior ability to recover biological signals missed by the standard analysis workflow in simulated ground truth datasets of cell type mixtures. When applied to a dataset of human multiple myeloma precursor conditions, HiDDEN recapitulates the expert manual annotation and discovers malignancy in previously considered healthy early stage samples. When applied to a mouse model of demyelination, HiDDEN identifies an endothelial subpopulation playing a role in early stage blood-brain barrier dysfunction. We anticipate that HiDDEN should find a wide usage in contexts which require the detection of subtle changes in cell types across conditions.

## Introduction

High-dimensional transcriptional profiling of cells has enabled the comprehensive characterization of cellular changes in response to perturbations, such as disease^1–4^, treatment with a drug^5^, or gene knockouts^6,7^. Existing computational strategies address different aspects of this general question, each accompanied by a set of assumptions^8^. Differential expression and differential abundance approaches aim to identify changes in gene expression and cell type proportion between perturbation conditions with the caveat that their power to infer the biological alterations is compromised when the condition-labels do not correctly represent the presence or absence of an effect in individual cells. For example, many perturbations only affect a subset of the cells in a given cell type while the rest of the cells are largely unaffected^9^. Condition-agnostic approaches aim to identify perturbation-affected groups of neighboring cells within the latent space, which may be clouded by the presence of several alternative axes of biological or technical variation^10–12^ making it challenging to tease out the perturbation-relevant signal.

To address these challenges, we developed a novel machine learning method called HiDDEN, which refines the labels of individual cells within perturbation conditions to accurately reflect their status as affected or unaffected. We systematically generate ground truth datasets of cell type mixtures and demonstrate that HiDDEN can find hard-to-detect affected subpopulations of cells and accurately identify their marker genes, outperforming the standard approach. We used HiDDEN to recapitulate manual annotation of neoplastic cells in human multiple myeloma precursor conditions and discover malignancy in previously considered healthy early stage samples, as well as to identify an endothelial cell subpopulation that regulates blood-brain barrier function during the early stages of demyelination in a mouse model.

## Results

### Overview of problem and method

In many case-control experiments, only a subset of the cells in case samples are affected by the perturbation (**Figure 1A**). The standard analysis workflow of jointly clustering gene expression profiles of case and control cells can fail to distinguish affected from unaffected cells, resulting in mixed clusters (**Figure 1B**) due to multiple sources of variance competing with the perturbation signal. Differential expression using the sample-level labels within a mixed cluster can fail to recover the perturbation markers due to the incorrect labels decreasing detection power (**Figure 1C**).

**Figure 1:**
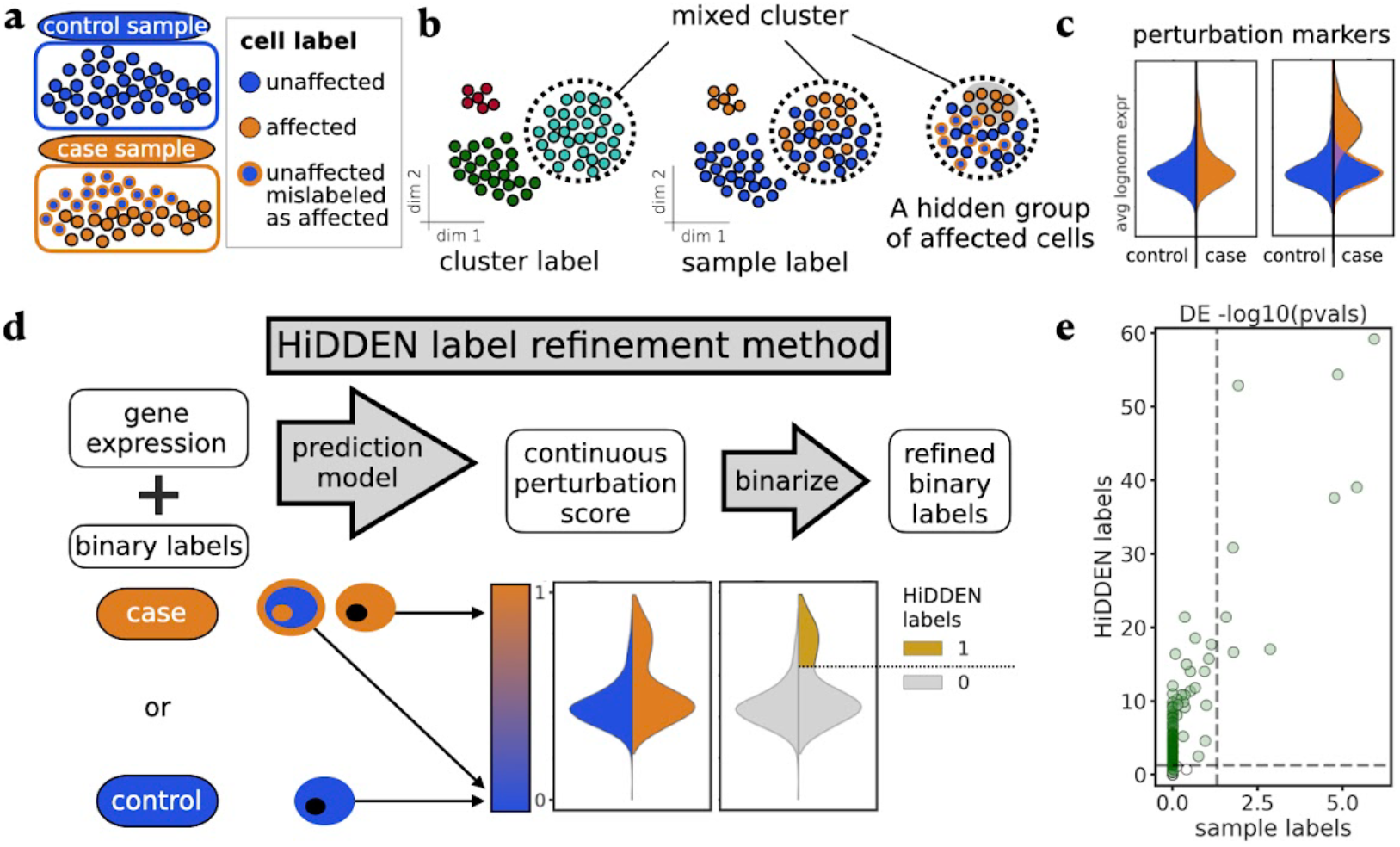
Overview of problem and HiDDEN label refinement method. **A** Setup of a case-control single cell experiment, in which all the cells in control samples are labeled as unaffected, while cells in case samples can be either affected or unaffected by the perturbation. **B** Standard clustering can produce clusters containing cells with mixed case-control sample-level labels while the subset of truly affected cells can be hidden. Colors as defined in A. **C** Representative violin plots of the average log normalized expression of perturbation markers split by sample-level labels (left) and highlighting the difference in the distributions of affected and unaffected cells within the case sample (right). Colors as defined in A. Area not scaled to count. **D** Overview of the HiDDEN label refinement method. First, a prediction model takes the gene expression profiles and the sample-level binary labels and transforms them into per cell continuous perturbation scores. Then, the continuous scores of cells originating from the case samples can be binarized into HiDDEN-refined binary labels (**Methods**). **E** Representative scatterplot of -log10 adjusted p-values per gene computed using differential expression (DE) on case-control sample labels (x-axis) and HiDDEN-refined binary labels (y-axis). Horizontal and vertical dashed lines drawn at -log10(0.05) significance threshold. Ground truth DE genes colored in green. Standard DE analysis on case-control labels misses ground truth markers, while HiDDEN successfully recovers many of them.

The standard analysis of single cell data is not tailored to identifying perturbation-associated signals. However, combining gene expression profiles and sample-level labels in a novel way allows us to leverage that at least some of the labels are correct and empowers HiDDEN to utilize the shared variability in features corresponding to correctly labeled cells. HiDDEN transforms the sample-level labels into cell-specific continuous perturbation-effect scores and assigns new binary cell labels, revealing their status as affected or unaffected (**Figure 1D, Methods**). The resulting binary labels can accurately capture the perturbation signature and boost power to detect genes whose expression is affected by the perturbation (**Figure 1E**).

### HiDDEN detects biological signal missed by the standard analysis workflow in simulated ground truth datasets of cell type mixtures

To simulate the biological change in cell function induced by a perturbation, we conducted simulations using the single-cell RNAseq profiles of Naive B and Memory B cells from a dataset of peripheral blood mononuclear cells (PBMC)^13^ (**Figure 2A, Methods**). Naive B and Memory B cells have relatively similar expression profiles but with biologically-relevant differences^14,15^, making them suitable for modeling perturbation-induced changes. To mimic the outcome of a perturbation experiment, we constructed a control sample consisting of Naive B (representing unperturbed) cells and a case sample consisting of both Naive B and Memory B (representing perturbed) cells (**Figure 2B, Methods**). We observed that, as Memory B and Naive B cells became increasingly imbalanced, the ability of a commonly used single cell analysis pipeline (**Methods**) to identify the Memory B cluster became impaired. For example, having 5% Memory B cells in the case condition results in a highly heterogeneous latent space produced by the standard dimensionality reduction workflow, making it impossible to detect a locus of perturbed cells (**Figure 2C**). Indeed, using a standard clustering workflow with default parameter values (**Methods**) fails to recover a cluster that purely represents the Memory B labeled cells (**Figure 2D**). Exposing the ground truth labels reveals that even the cluster with the highest enrichment of case-labeled cells contains a majority of Naive B cells (**Figure 2D**). The recovery of the Memory B cluster could not be improved by varying the number of principal components (PCs) used to construct the latent space, adjusting the resolution parameter of the clustering algorithm, or varying the gene selection, including by utilizing Naive B and Memory B markers in lieu of highly variable genes (**Supplementary Figure 1**). By contrast, HiDDEN was far better able to identify the Memory B signature within this artificial mixture and distinguish Memory B from Naive B cells (**Figure 2E**).

**Figure 2:**
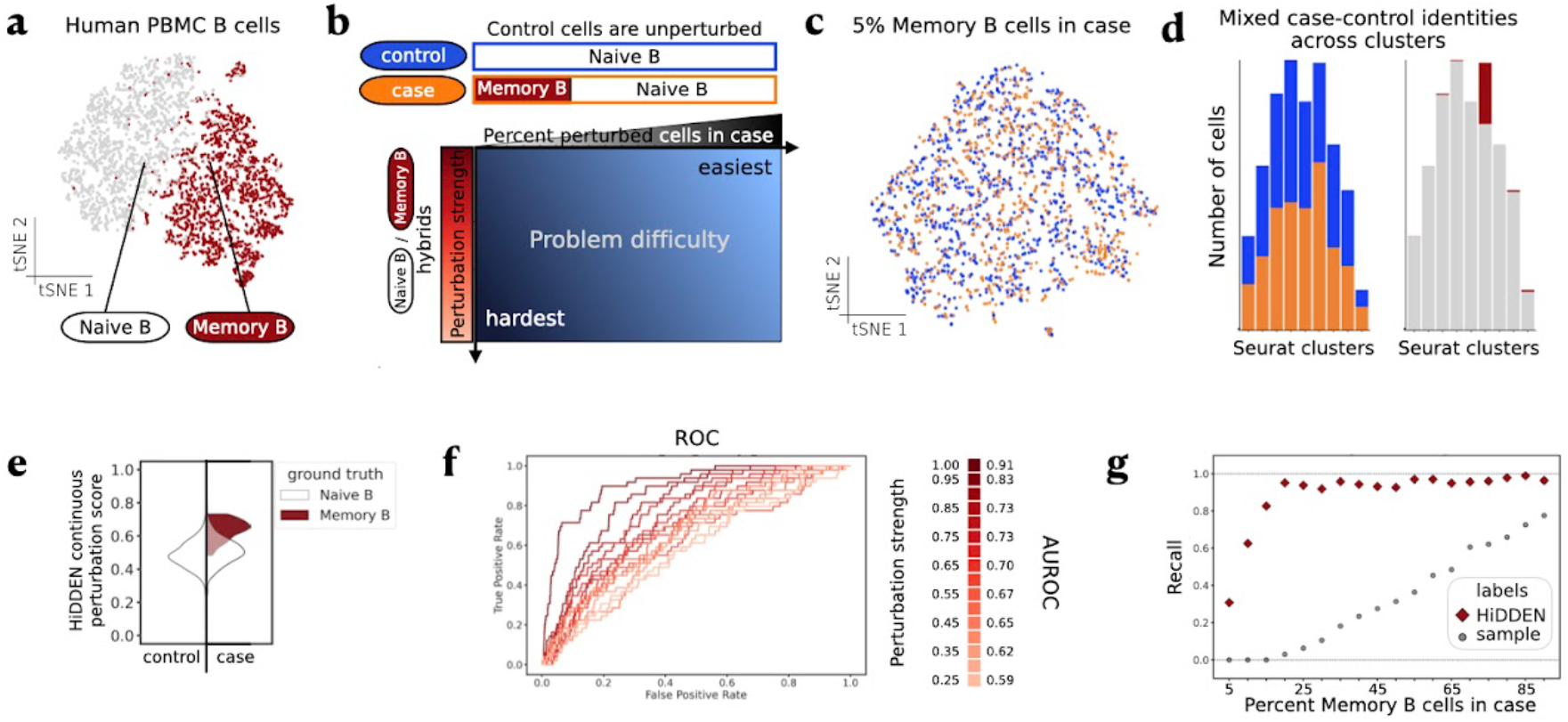
HiDDEN detects biological signal missed by the standard analysis workflow in simulated ground truth mixtures of two cell types. **A** tSNE embeddings of Naive B and Memory B cell gene expression. **B** Schematic of problem difficulty and definition of synthetic datasets along two axes: percent perturbed cells in case sample (x-axis) and strength of the perturbation (y-axis). Detecting the perturbation is most challenging when there are few affected cells and the difference between affected and unaffected cells is small. **C** tSNE embeddings of a representative simulated dataset containing 5% Memory B cells in the case sample. Cells colored by case-control labels. Colors as defined in B. **D** Distribution of case-control (left) and Memory B-Naive B (right) cell identities across Seurat clusters. Colors as defined in A and B. **E** Violin plot of the distributions of the continuous perturbation score of Naive B and Memory B cells split over control and case and colored by ground truth labels, for the dataset containing 5% Memory B cells in the case sample. **F** Area under the Receiver Operating Characteristic (ROC) curves for classification of ground truth cell labels as a function of percent perturbed cells in case sample with the Area Under the ROC (AUROC) indicated in the legend for a sampling of the curves, for the dataset containing 5% perturbed cells in the case sample. **G** Recall of ground truth DE genes by DE testing on indicated labels as a function of percent perturbed cells in case sample.

This example reveals a more general feature of how cell types are detected in single cell data. To comprehensively characterize the problem difficulty and assess the power of our method to detect the perturbation signal, we constructed a collection of ground truth case-control datasets by varying two key aspects (**Figure 2B, Methods**). First, to study the effect of perturbation strength, we defined perturbed cells as hybrids of Naive B and Memory B cells of variable relative weight (**Methods**). Decreasing the strength of the transcriptional difference between perturbed and unperturbed cells increased the difficulty of identifying the perturbed cell cluster (**Figure 2F, Supplementary Figure 2, Methods**). Even when only 5% of the case cells are Memory B, HiDDEN continuous perturbation scores identified the biological differences between Memory B and Naive B cells with high accuracy. Second, to explore the influence of class imbalance, we varied the percent of perturbed cells in the case sample (**Supplementary Table 1**). Strikingly, in datasets with fewer than 20% Memory B cells, using the standard analysis pipeline with the sample-level labels completely failed to retrieve any of the Naive B and Memory B marker genes, and overall retrieved only a fraction even in datasets with high sample label accuracy (**Figure 2G, Methods**). The marker gene recovery was not improved even when we considered the union of case-control label derived DE genes per cluster since the reduction in number of cells dramatically hinders the power of DE testing to recapitulate the markers (**Supplementary Figure 3, Methods**). By contrast, the HiDDEN-refined binary labels had superior power to detect the ground truth markers. Furthermore, HiDDEN-refined labels appeared to provide accuracy beyond the ground truth labels for this dataset. Specifically, the genes identified by DE testing on HiDDEN-refined labels, but not by DE testing on ground truth labels, identified additional genes that are consistent with markers of Naive B and Memory B cells (**Supplementary Figure 4**), suggesting that HiDDEN labels possess corrective power for a slight amount of misclassification that might have occurred in the original annotation.

Several computational methods have recently been proposed to characterize perturbation effects in single-cell data, each designed to tackle a particular aspect of this general problem. Specifically, CNA^11^ provides cluster-free detection of perturbation-affected areas of the latent space, while MELD^16^ offers identification of a perturbation gradient. A third method, Milo^17^, performs differential abundance testing over continuous trajectories. When applied to our target task of refining the sample-level case status into perturbed and unperturbed cell labels, HiDDEN continuous perturbation scores and binary-refined labels outperformed the corresponding continuous and binarized scores from CNA, MELD, and Milo across ground truth Naive B / Memory B mixtures (**Supplementary Figure 5, Methods**). Each of these methods relies on veritable cell-level labels as input, such that HiDDEN-refined labels could augment their respective performances. Indeed, when HiDDEN is applied first, there was an improvement in performance of CNA, MELD, and Milo-derived continuous scores (**Supplementary Figure 6, Methods**) and binarized labels (**Supplementary Figure 7, Methods**) to recover the ground truth perturbation labels.

The HiDDEN method has a single model parameter: the number of features in the predictive model. To explore how this parameter affects the stability and accuracy of HiDDEN results, we generated two heuristics for automatically choosing it and demonstrated that either strategy works well, indicating that the parameter is not especially influential for model performance (**Supplementary Figure 8, Methods**).

### HiDDEN recapitulates manual annotation of neoplastic cells in human multiple myeloma precursor conditions and discovers malignancy in previously considered healthy early stage samples

To test the ability of our method to capture perturbation signal in a real dataset, we applied HiDDEN to single-cell RNA-seq profiles of human bone marrow plasma cells from patients with multiple myeloma (MM), its precursor conditions smoldering multiple myeloma (SMM) and monoclonal gammopathy of undetermined significance (MGUS), and healthy donors with normal bone marrow (NBM)^4^ (**Methods**). Precursor samples can contain a mixture of neoplastic and normal cells (**Figure 3A**) and the authors of the original study defined two orthogonal strategies for describing the malignancy status of precursor samples and their cells. The first strategy is a per-sample computational analysis excluding immunoglobulin light chain genes followed by manual annotation resulting in binary labels defining cells as healthy or malignant. The second strategy is a tumor-purity estimate of the proportion of malignant cells in each precursor sample from a model based on the distribution of immunoglobulin gene expression.

**Figure 3:**
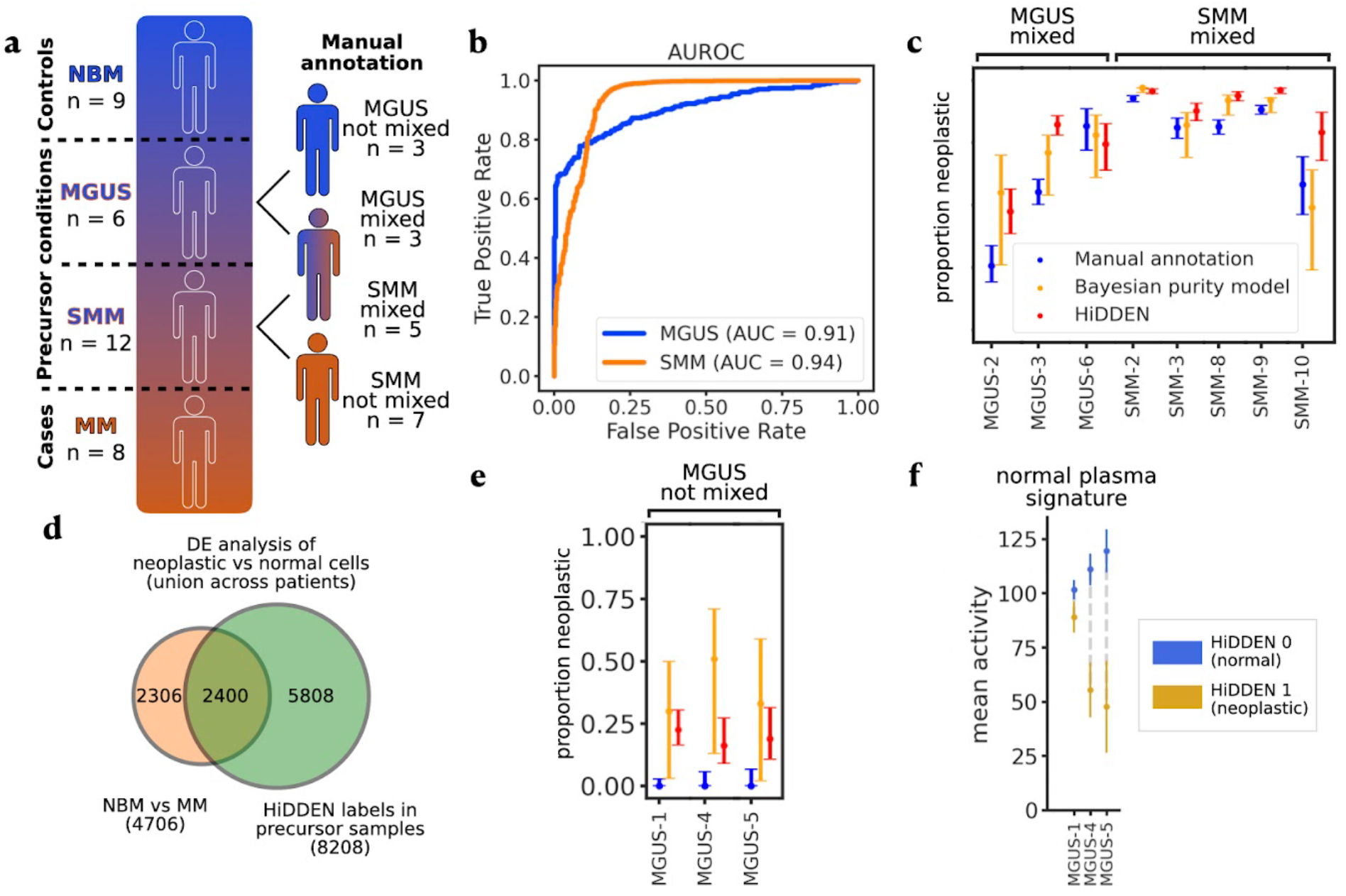
Application of HiDDEN to a human bone marrow dataset with previously published annotations. **A** Schematic of human bone marrow dataset^4^ which includes human plasma cells from healthy donors, multiple myeloma patients, and two precursor states. Precursor samples possibly contain a mixture of healthy and malignant cells. Manual annotation of healthy and malignant cells per precursor patient were reported previously^4^. **B** AUROC for predicting the per-cell malignancy status in mixed samples averaged for each precursor state. **C** Comparison of manual annotation, Bayesian purity model, and HiDDEN predictions for estimating the per-sample neoplastic proportion (y-axis) in mixed MGUS and SMM samples (x-axis). **D** Venn diagram of DE genes comparing neoplastic with normal cells based on the NBM/MM samples and the HiDDEN refined labels in precursor samples identified 2400 significantly overlapping genes (hypergeometric test, p-value=3.066e-31) and 5808 genes uniquely found using HiDDEN labels in precursor samples. **E** Comparison of manual annotation, Bayesian purity model, and HiDDEN predictions for estimating the per-sample neoplastic proportion (y-axis) in non-mixed MGUS samples (x-axis). Colors as defined in C. **F** Computational validation of cells predicted to be malignant by HiDDEN in low tumor purity MGUS samples. Mean activity (y-axis) and confidence bounds of genes assigned to a biologically-interpretable normal plasma signature from the original study separately computed per HiDDEN label in each low tumor purity MGUS sample (x-axis).

According to the manual annotation, three MGUS and five SMM samples contain a mixture of malignant and healthy cells. Application of HiDDEN continuous perturbation scores to distinguish cells in these mixed samples showed remarkable agreement with this manual annotation (**Figure 3B, Supplementary Figure 9, Methods**). Furthermore, sample purity estimates derived from HiDDEN binary labels agreed with their point estimates of malignant cell proportions, outperforming manual annotation-based estimates in the majority of mixed precursor samples (**Figure 3C, Methods**).

We next turned our attention to the three MGUS samples with the lowest tumor purity. In those samples, the manual annotation strategy failed to identify any neoplastic cells. By contrast, HiDDEN was able to discover malignant cells in these early stage patients that were missed by the manual annotation (**Figure 3E, Methods, Supplementary Table 2**). To computationally validate that these were indeed malignant cells, we assessed whether the distinguishing genes of these cells matched with known signatures of healthy plasma and malignancy. Indeed, previously described gene signatures distinguishing normal from malignant cells^4^ were heavily differentially enriched between the HiDDEN-defined normal and malignant cells in these samples (**Figure 3F, Supplementary Figure 10, Methods**).

Of note, HiDDEN successfully recapitulates both types of malignancy estimates in the presence of pronounced patient-specific batch effects in this dataset (**Supplementary Figure 11A**). Mirroring the analysis in the original study, we deployed a batch-sensitive strategy to fitting the HiDDEN model, namely training it on all NBM, all MM, and one precursor sample at a time. Additionally, we also developed a batch-agnostic strategy, where we fit all samples together (**Methods**). HiDDEN outputs under both strategies were closely aligned and almost indistinguishable (**Supplementary Figure 11B-F**). We provide a heuristic to automatically choose the optimal number of features used in the training of the prediction model and demonstrate that HiDDEN outputs were closely aligned and almost indistinguishable across a wide range of values for the tunable model parameter (**Supplementary Figure 12, Methods**).

We leveraged our refined definition of healthy and neoplastic cells in precursor states to derive markers of early disease. We find a total of 8208 differentially expressed genes, 2400 of which significantly overlap (hypergeometric test, p-value=3.066e-31) with basic malignancy markers derived from a comparison of healthy and multiple myeloma patients, and 5808 of which are uniquely found using the HiDDEN-refined labels in precursor samples (**Figure 3D, Methods**).

### HiDDEN identifies an endothelial subpopulation affected in the early stages of demyelination

To explore HiDDEN’s ability to identify rare, subtle perturbations, we applied the method to single-nucleus RNA-seq (snRNA-seq) profiles from a time-resolved dataset of a mouse model of demyelination^18,19^ (**Figure 4A, Methods**). In this experiment, case animals received a corpus callosum injection containing lysophosphatidylcholine (LPC), a compound toxic to oligodendrocytes, while control animals are injected with saline (PBS) (**Methods**). LPC induces white matter loss, demyelination, which is rapidly repaired in a stereotyped manner over three weeks. Several cell types showed dramatic changes in response to this injury (**Supplementary Figure 13A**) as expected^20–22^, but the effects on endothelial cells (ECs) appeared modest (**Figure 4B,C**). As vascular cells of the brain, ECs play critical roles in homeostasis, myelin formation and tissue repair, but the altered genes and pathways underlying these functions in demyelination are poorly understood^23,24^.

**Figure 4:**
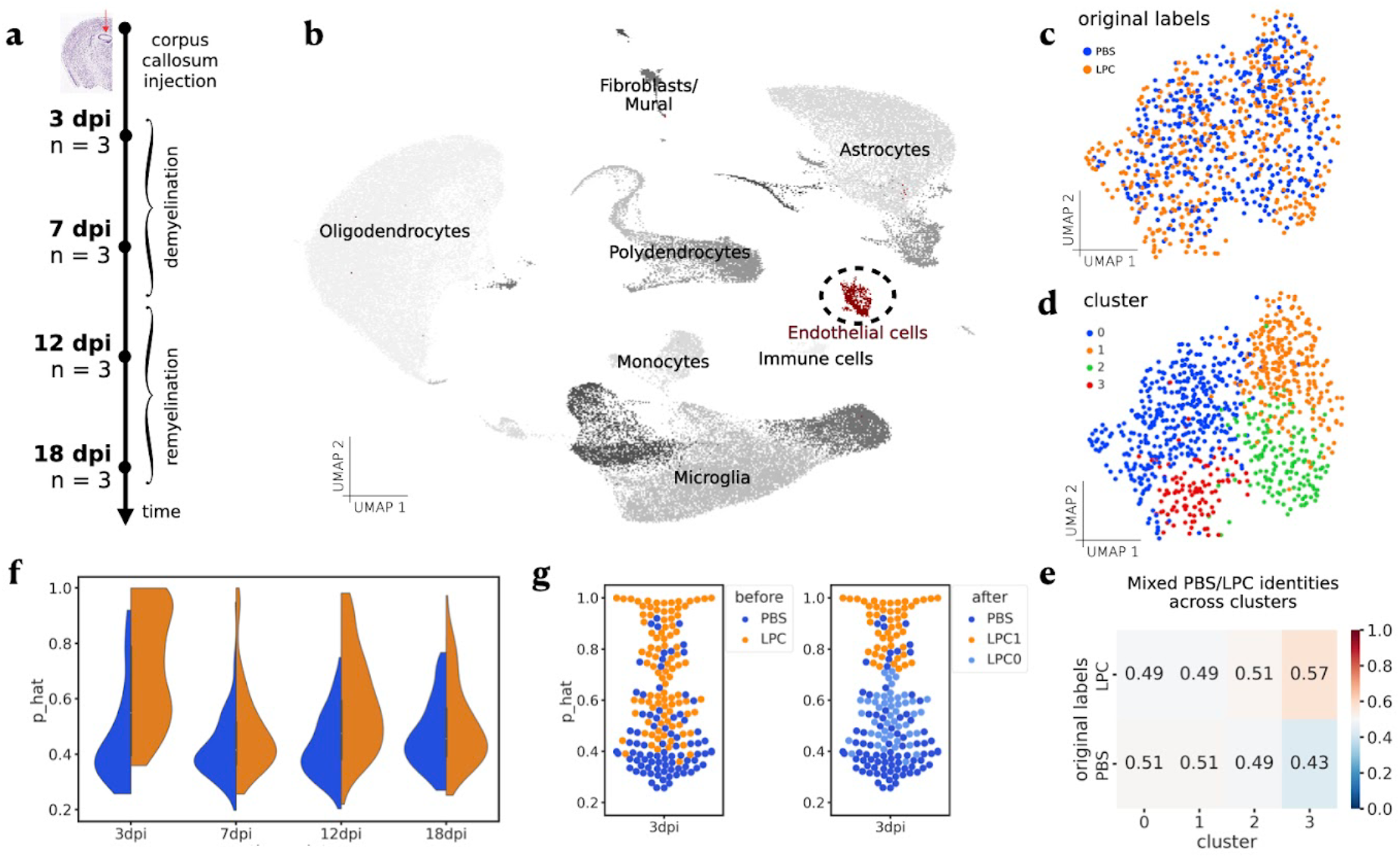
Application of HiDDEN to ECs from a mouse demyelination time-course experiment. **A** Overview of experimental design. Corpus callosum injection with saline (PBS) and a compound toxic to oligodendrocytes (LPC) used to induce demyelination with n=3 mice per condition per time point across four time points. **B** UMAP embeddings of non-neuronal cells from PBS (control) and LPC conditions across all time points colored by annotation of major cell type. ECs highlighted in red. UMAP embeddings of ECs across all time points colored by PBS/LPC sample-level labels (**C**), and Seurat cluster labels (**D**). **E** Relative abundance of case-control cell identities across Seurat clusters. **F** Violin plots of HiDDEN continuous perturbation scores split over PBS and LPC labels and grouped by time point. **G** Swarmplot of HiDDEN continuous perturbation scores for the 3dpi cells colored by original PBS/LPC labels (left) and with color indicating the refinement of LPC cells into affected (LPC1) and unaffected (LPC0) (right).

We first examine ECs during remyelination using an existing analytic pipeline^25^. The standard dimensionality reduction workflow produced a homogeneous distribution of sample-level labels in the latent space (**Figure 4C, Methods**) and clustering failed to identify a perturbation-enriched subpopulation (**Figure 4D, E, Methods**). The case-control identities were similarly mixed across time points (**Supplementary Figure 13C**). By contrast, fitting HiDDEN to the ECs across all time points (**Methods**) generated a bimodal distribution of continuous perturbation scores for case cells at the earliest time point, suggesting an underlying mixture of affected and unaffected cells (**Figure 4F, Supplementary Figure 13D**). We split the bimodal distribution of continuous scores and used the resulting binary cell-labels to define demyelination-affected and unaffected EC subpopulations, denoted LPC1 and LPC0, respectively (**Figure 4G, Methods**). Together this demonstrates that HiDDEN can reveal demyelination-specific effects on endothelial cells not apparent using conventional analysis approaches.

We next analyzed the differential response of LPC1 and LPC0 ECs to demyelination. The LPC1 subpopulation was characterized by 28 unique markers (**Figure 5A, Supplementary Table 3, Methods**), a subset of which we experimentally validated to be lesion-specific with *in-situ* hybridization at the 3dpi timepoint (**Figure 5B**). To understand the biological functions of these changes, we applied Gene Set Enrichment Analysis. This revealed the observed changes in gene expression were consistent with alterations that occur in the context of inflammation and demyelination, such as increased angiogenesis ^26^, blood-brain barrier breakdown^27,28^, and increased production of extracellular matrix components (**Figure 5C, D, Methods**). Together, this suggests that the LPC1 EC subset, revealed by the HiDDEN method, have altered core endothelial functions specifically during the early stages of white matter damage.

**Figure 5:**
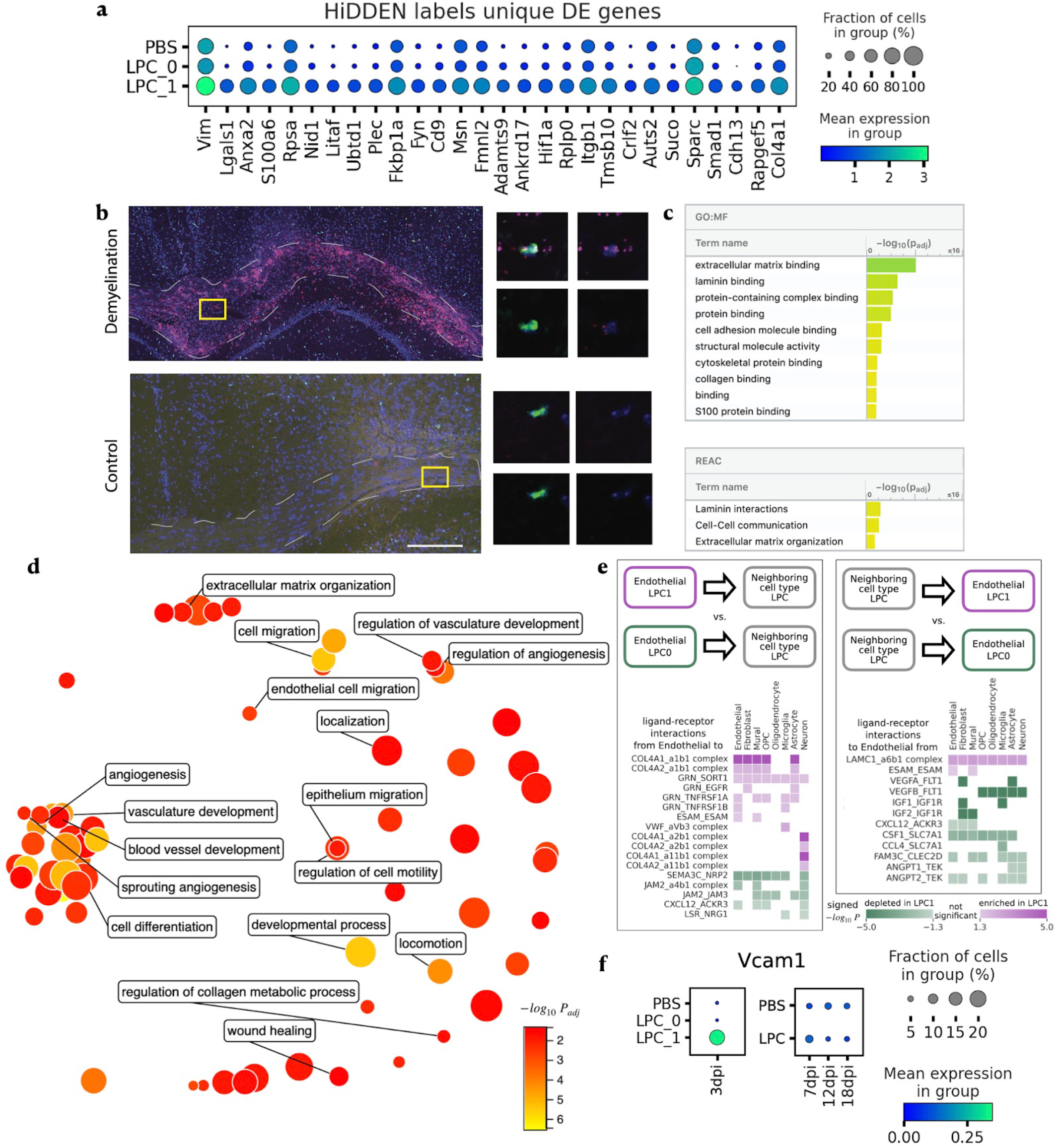
Characterization of the demyelination-affected endothelial subpopulation (LPC1) identified by HiDDEN. **A** Dotplot of mean expression of LPC1 marker genes ordered by p-value. **B** Validation of gene expression with fluorescent in-situ hybridization (FISH) and confocal microscopy showing presence of endothelial cells (*Flt1*-positive cells, green) coexpressing *Lgals1* (red) and *S100a6* (magenta) specifically present in demyelinating lesion (top) and not control (bottom) brains, 3 days after injection. Left: Overview of a demyelinating or control white matter lesion. Corpus callosum outlined in gray dashed line. Right: High-resolution confocal images of single endothelial cells. All images are representative of n=2-3. **C** Significantly enriched GO molecular function terms (top) and Reactome pathways (bottom) based on LPC1 marker genes ordered by significance. **D** ReviGO plot summarizing the significantly enriched GO biological processes based on LPC1 marker genes colored by significance with selected labels. **E** Significantly enriched (purple) and depleted (green) ligand-receptor interactions between LPC1 (relative to LPC0) endothelial cells and neighboring cell types split by interaction direction: from endothelial to neighboring cell type (left), and from neighboring cell type to endothelial (right). **F** Dot plots of mean expression of Vcam1 across time and condition labels.

The blood-brain barrier is an active hub for cell-cell interactions between the ECs comprising blood vessel walls and the surrounding cell types. In particular, endothelial-endothelial, endothelial-fibroblast, and endothelial-astrocyte interactions are crucial in tightly regulating blood-brain barrier permeability^29^ and have been implicated in neurodegenerative disease pathogenesis^28^. To investigate if these intercellular pathways could be dysregulated during a demyelinating event, we examined differences in cellular communication of the affected LPC1 and unaffected LPC0 endothelial subpopulations with neighboring cell types, we used a computational method for targeted hypothesis testing of ligand-receptor expression (**Figure 5E, Supplementary Figure 14, Methods**). The changes in communication were consistent with increased angiogenesis, blood-brain barrier breakdown, and increased extracellular matrix. The anti-angiogenic interactions of *Flt1* with *Vegfa*/*Vegfb* and of *Sema3a* with *Npr2* were decreased in LPC1 endothelial cells, supporting increased angiogenesis. In addition, interactions between collagen and integrin components were increased in LPC1, pointing to remodeling of the extracellular matrix. Interactions supporting the tight junctions between endothelial cells, such as *Jam2* and *Jam3* with integrins, were also decreased in LPC1, suggesting compromised barrier function. Furthermore, we found that LPC1 endothelial cells have an increased expression of *Vcam1*, which acts as a ligand for recruiting immune cells from the bloodstream to cross the blood-brain barrier (**Figure 5F**). In summary, changes in EC function during white matter damage are poorly understood. While established approaches failed to identify changes in ECs during de- and remyelination, HiDDEN revealed a temporally-specific alteration of a subset of ECs during demyelination which likely drives blood-brain barrier breakdown and immune cell influx. As few biological contexts or perturbations are truly uniform, this illustrates the power and broad utility of HiDDEN to isolate *bonafide* effects from complex biological systems *in vivo*.

## Discussion

In this work, we introduced a novel computational method, HiDDEN, which accurately refines the condition labels of case cells into affected and unaffected for more sensitive detection of perturbation signals. We leveraged the HiDDEN output to find hard-to-detect disease-affected subpopulations of cells and characterized their marker genes using differential expression testing. We provide a computationally efficient Python implementation of HiDDEN at https://github.com/tudaga/LabelCorrection, making it scalable to large datasets. At the same time, in the application to endothelial cells, we found that HiDDEN is capable of detecting subtle perturbation changes involving tens of genes even in small datasets on the order of a hundred cells. In the application to human bone marrow plasma cells, we showed that HiDDEN can be successfully applied to samples from heterogeneous conditions with pronounced batch effects without the necessity of sophisticated preprocessing or alignment. Therefore, HiDDEN is highly applicable to single-cell atlases with batch effects and a high variability in retrieved numbers of cells across cell types.

Detecting cell-level transcriptional changes across experimental conditions is one of the big promises of high-resolution single-cell expression data^8^. In recent years, several methods have been proposed to characterize perturbation effects in single-cell data. The standard analysis workflow performs label-agnostic dimensionality reduction and clustering, followed by comparisons of cell attributes across condition labels within clusters. CNA^11^ provides a cluster-free approach identifying regions in the latent space of uneven mixing of condition labels. MELD^16^ produces a continuous measure of the perturbation effect by distributing the condition labels among neighbors in the cell state manifold. Milo^17^ performs differential abundance testing among experimental conditions in the presence of continuous trajectories. Mixscape^30^ removes known confounding sources of variation and dissects successfully from unsuccessfully perturbed cells. These approaches rely on at least one of the following assumptions: 1) the condition labels correctly represent the presence or absence of an effect in individual cells; 2) the perturbation effect is a dominant signal in the latent space; or 3) that any confounding sources of variation are known and can be removed. However, these assumptions might not always be met – often, perturbation effects are outshined by biological heterogeneity and technical noise or the affected subpopulation of cells is small and therefore the condition labels are mostly incorrect. We demonstrate that the HiDDEN-refined binary labels can be used to boost the performance of existing approaches relying on cell-level labels accurately representing the presence of a perturbation effect, including CNA and MELD, as well as methods for differential abundance across conditions, such as Milo.

In this paper, we focused on applications of HiDDEN to detect the presence or absence of a disease effect at single-cell resolution. However, our method is versatile and can be applied to any context in which the aim is to focus the latent space on a particular distinction, for example to explore subtle genotype effects (i.e. eQTLs) and sexual dimorphism. The HiDDEN framework is amenable to extensions to spatial and multi-omics data, as well as applications beyond a binary output, such as multi-stage disease progressions or time-course experiments.

## Methods

### HiDDEN: A computational method for revealing subtle transcriptional heterogeneity and perturbation markers in case-control studies

#### Intuition

In a case-control experiment, typically all of the cells in control samples will be unaffected, and possibly only a subset of the cells in case samples will be affected by the perturbation (**Figure 1A**). Using the gene expression profiles alone can fail to separate out the affected from unaffected cells (**Figure 1B**). Using the sample-level labels alone can fail to recover the perturbation markers (**Figure 1C**). However, combining the two allows us to leverage that at least some of the labels are correct and allows a prediction model to utilize the shared variability in features corresponding to correctly labeled cells. As a result, we transform the sample-level labels into cell-specific perturbation effect scores and can assign binary cell labels representing their status as affected or unaffected (**Figure 1D**). We use this information to find hard-to-detect affected subpopulations of cells, characterize their marker genes, and contrast their cellular communication patterns with those of unaffected cells.

#### Notation

Let *X* ∈ *R*^*N* × *M*^ denote the matrix containing the gene expression profiles of *N* cells across *M* genes. Let *Z* ∈ *R*^*N*×*M*^ denote the reduced representation of the *N* cells in a *K*-dimensional latent space of features. Let *Y* ∈ {0,1}^*N*^ denote the binary vector encoding the sample-level label of each cell, where 0 stands for control and 1 stands for case. We train a predictive model denoted by *h*(.) on the reduced representation *Z* and the binary sample-level labels *Y*. Due to the binary nature of the case-control labels *Y*, the predictive model is a binary classifier modeling the probability of label 1 given the input features, i.e. *P*(*Y* = 1|*Z*) = *h*(*Z*). We train the parameters of the classifier on a dataset of interest and denote the fitted value of *P*(*Y* = 1|*Z*) with 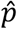. Finally, we cluster the continuous scores 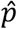 to derive refined binary labels, denoted by 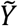, reflecting the status of each cell as affected or unaffected by the perturbation regardless of which sample it originated from.

#### Construction of the latent space

We transform the *N* × *M* gene expression matrix *X* into an *N* × *K* matrix *Z* containing an information-rich reduced representation of the gene expression profile of each cell in the dataset. Throughout this work we use principal component analysis (PCA) for this task, which is a commonly used dimensionality reduction technique for single-cell RNA-seq data. In principle, any other information-preserving dimensionality reduction method can be used to construct the latent features in lieu of PCA, such as non-negative matrix factorization (NMF), or the latent representations from an autoencoder. We did not find convincing evidence that using more sophisticated dimensionality reduction techniques affected model performance (data not shown) and thus adopted PCA as the most computationally efficient option.

#### Estimation of the continuous perturbation score

For each cell, we derive a continuous score, 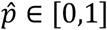, reflective of the strength of the perturbation effect on that cell relative to the rest of the cells in the dataset. Throughout this work we use logistic regression for this task. That is, the predicted probability of label 1 given the input features is given by

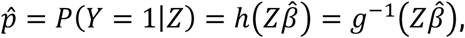

where *h* is the logit link function, i.e. *g* is the logistic function. In principle, we can use any classifier *h*: *R*^*N*×*M*^ → [0,1]^*N*^, including large-parameter non-linear models such as neural networks. In practice, we opted for logistic regression as a simple yet powerful model with a canonical parameter optimization routine that does not introduce additional hyperparameters, training heuristics, and increased computational resources and time demands.

#### Derivation of the refined binary label

Each cell in a dataset from a case-control experiment possesses a binary sample-level label reflecting whether it originated from a case or control sample. As these coarse labels do not reflect the individual cell identity of being affected or unaffected by the perturbation, we derive a new refined binary label that captures the presence or absence of a perturbation effect in each cell. When we want to distinguish between affected and unaffected cells in the case sample, we cluster the continuous perturbation scores 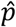 for all the cells with initial label 1 into two groups. The cells in the group with lower 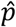 scores receive new label 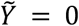 and the cells in the group with higher 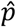 scores get a new label 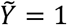 matching their old label. To do this, we use k-means clustering with *k* = 2. In principle, any other clustering algorithm, such as Gaussian mixture models for example, can be utilized in lieu of k-means. However, in practice the choice of clustering method is not consequential, and they tend to perform equivalently since we are clustering a 1-dimensional vector into two groups.

#### Choosing the number of latent dimensions

When selecting *P*, the number of latent dimensions, our guiding principle is that *P* should be chosen in a data-dependent manner aiming to retain informative transcriptional heterogeneity while avoiding overfitting the not-entirely-correct sample-level labels. For example, note that using *P* > *rank*(*X*) will yield predictions 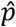 synonymous with the sample-level labels. In practice, this implies we should choose *P* << *min*(*N, M*). Therefore, we develop two novel data-driven heuristics to quantify the amount of informative heterogeneity retained in the latent space by measuring the perturbation signal downstream of redefining the binary labels.

The first heuristic is to use the number of differentially expressed genes defined by the refined labels 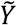. A large number of DE genes indicates a meaningful signal in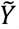, and in turn in 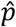 and the latent space *Z*. Intuitively, when *P* is too small, the continuous scores 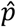 do not contain enough heterogeneity to yield labels 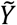 that distinguish DE genes. Conversely, when *P* is too large, we are overfitting to the sample-level labels resulting in low power to detect the perturbation markers. To achieve an appropriate balance, we scan a range of values for *P* that traverses the concave relationship between *P* and the number of DE genes and choose the number of latent dimensions maximizing it.

The second heuristic is to use the strength of the difference between the values of 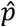 for cells in the case sample with new label 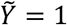 and new label 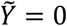. We quantify the probability that these two sets of values are drawn from the same distribution using the two-sample Kolmogorov-Smirnov (KS) test. The larger the value of the KS test statistic, the more different the sample distributions of the perturbation score are between the cells predicted to be affected and unaffected. This value is generally an increasing function of *P*, therefore we pick the smallest value of *P* that maximizes the KS test statistic.

Note that we do not need access to ground truth labels of the perturbation effect for neither heuristic. When ground truth data is available, we find that the ability of the refined binary labels 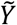 to represent the true perturbation effect per cell is similarly high for a wide range of values of *P* and that either heuristic yields a choice in that range.

### Assessing performance on semi-simulated ground truth data

To demonstrate and quantify the problem difficulty and to assess the power of our method, we conducted simulations using a real single-cell dataset. We used the RNA profiles of *n* = 1900 Naive B and *n* = 1630 Memory B cells from a dataset of peripheral blood mononuclear cells (PBMC) freely available from 10x Genomics^13^. We first describe our design of simulated case-control datasets by combining the two B cell subpopulations. We then describe the challenge of separating the two subpopulations using the standard single cell clustering analysis workflow. Finally, we describe how we train our method and the metrics we use to assess its power to detect the biological signal.

#### Generation of ground truth datasets

The RNA profiles of all *n* = 30672 cells in the human PBMC data from 10x Genomics were clustered and annotated independently of the ATAC-seq profiles following standard approaches detailed in the Seurat Weighted Nearest Neighbor Analysis vignette^31^. This resulted in 27 annotated cell types of which we subset all Naive B and Memory B cells for the subsequent generation of ground truth datasets.

For the tSNE representation of all Naive B and all Memory B cells in **Figure 2A**, we normalized the gene counts, performed variable gene selection, scaled the normalized counts, performed dimensionality reduction using PCA, built the nearest-neighbor graph, and ran tSNE, all with default hyperparameter values using the standard functions in Seurat v 3.2.3^25^.

To comprehensively describe the problem difficulty and test the performance of our method, we constructed a collection of ground truth case-control datasets by varying two aspects (**Figure 2B**). In each dataset, the control sample consists entirely of Naive B cells, which we refer to as unperturbed, whereas the case dataset consists of both unperturbed and perturbed cells, which are either Memory B or hybrids of Naive B and Memory B cells. Each dataset is indexed by (1) the percent perturbed cells in the case sample and (2) the strength of the perturbation.

To explore the effect of the percent perturbed cells in the case sample, we randomly drew (100 − *p*) % of the cells in the case from the Naive B cells and *p* % from the Memory B cells. The remaining Naive B cells were all allocated to the control sample. We explored 18 values of the percent perturbed cells *p* ∈ {5, 10, 15, …, 85, 90} %. The resulting number of Naive B and Memory B cells across case and control per dataset is provided in **Supplementary Table 1**.

To explore the effect of the strength of the perturbation, we varied the extent to which perturbed and unperturbed cells in the case sample differ from each other. Let *W*^(*m*)^ and *W*^(*n*)^ denote the weight of Memory B and Naive B contribution, respectively, for each hybrid cell, where *W*^(*m*)^ + *W*^(*n*)^ = 1, and *W*^(*m*)^, *W*^(*m*)^ ≥ 0, i.e. *W*^{*m*} denotes the strength of the perturbation. Let *i* index the perturbed (Memory B) cells in the case sample and let *N*, denote the total number of UMIs in cell *i*. Each memory B / Naive B hybrid cell has the same number of UMIs, *N*,, as the Memory B cell it originates from. Let 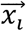 denote the gene expression profile of the original Memory B cell *i*. Let 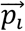 denote the normalized counts, i.e. the relative proportion of counts across genes. We drew a Memory B / Naive B hybrid gene expression profile 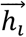 by first subsampling 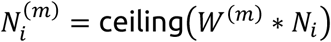 counts from the original Memory B expression profile 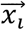:

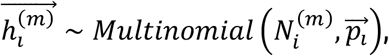

then we drew 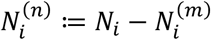 counts from the Naive B centroid, 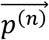, defined as the average normalized counts vector across all Naive B cells in the dataset:

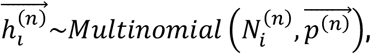

and finally, summed the two count vectors 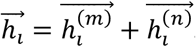.

We explored 17 values of the perturbation strength parameter *W*^(*m*)^ ∈ {0.25, 0.3, 0.35, …, 0.95, 1}. Overall, spanning both the percent perturbed cells in case and the perturbation strength axes, we generated 72 datasets to characterize the problem difficulty and assess the performance of our method, as described below.

#### Clustering analysis of ground truth datasets

For the tSNE representation in **Figure 2C**, we focused on the simulated dataset containing 5% Memory B cells in the case. We used the standard Seurat functions and normalized the gene counts, performed variable gene selection with default parameter values, scaled the normalized counts, performed dimensionality reduction using PCA with default parameter values, built the nearest-neighbor graph with the default number of nearest neighbors, used the Leiden algorithm with the default value of the resolution parameter to find clusters, and ran tSNE. For the bar plots in **Figure 2D** and **Supplementary Figure 1**, we computed the abundance of case and control-labeled cells and the abundance of Naive B and Memory B cells in each cluster.

Several hyperparameters influence the clustering results, and we varied each one to study their effects on the problem difficulty. The challenge of capturing the biological signal and separating Memory B from Naive B cells using the standard pipeline lies in: (1) the construction of the latent space; and (2) the resolution parameter of the clustering algorithm. To quantify the problem difficulty, we investigated the degree of separability of Naive B and Memory B cells in the latent space via the distribution of the number of Memory B nearest neighbors across Memory B cells. For a given simulated dataset, varying the number of principal components (PCs) used to build the nearest-neighbor graph impacts the separability of the latent space with including more PCs resulting in a more mixed latent space (**Supplementary Figure 1**). The choice of a feature selection strategy along with the choice of the resolution parameter of the Leiden clustering algorithm have a significant impact on the results. We observed that using highly variable gene selection with the resolution parameter chosen to yield two clusters (since we are aiming to separate two cell types) fails to isolate the Memory B cells (**Supplementary Figure 1**). In this simulated ground truth setting, we can compute the differentially expressed (DE) genes, i.e. marker genes, between the two classes and use them as the selected features. However, that choice alone is also not sufficient to yield improved clustering when using the default value of the resolution parameter.

#### HiDDEN model training

The HiDDEN model training was done in python and consists of three steps: (1) we preprocess the raw gene expression counts, (2) we train a logistic regression model, and (3) we binarize the predictions for cells in the case sample. First, we followed the standard preprocessing routine for single-cell RNA-seq data in scanpy^25^ which consists of filtering out cells with < 20 genes and filtering out genes not expressed in any cells, followed by log-normalization, and then, we used the standard PCA dimensionality reduction routine in scanpy on the scaled gene features. Second, we used the LogisticRegression function from the sklearn.linear_model python library to train a logistic regression on the binary sample-level labels and the first *P* PCs. Finally, we used the KMeans function from the sklearn.cluster python library with *n_clusters* = 2 on the continuous perturbation scores output by the logistic regression for cells in the case sample.

Note that HiDDEN does not require parameter tuning. Since we use all genes when computing the PC embedding of the data, the only parameter in the model is *P*, the number of PCs used in the training of the logistic regression. As described earlier in this section, we provide two data-driven heuristics for automatically choosing an appropriate value for *P*. For each dataset and for each heuristic, we scanned all integer values for *P* in the range [2, 60]. The first heuristic is to choose *P* that maximizes the number of DE genes defined by the HiDDEN refined binary labels. To compute the number of DE genes downstream of a given value of *P* \*in* [2, 60], we used the Wilcoxon rank-sum differential expression test in scanpy with adjusted p-value threshold < 0.05 (**Supplementary figure 5**). The second heuristic is to choose the smallest value of *P* that maximizes the value of the two-sample Kolmogorov-Smirnov (KS) test statistic comparing the sampling distributions of 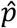 for cells in the case sample with new refined label 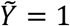 and 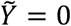 (**Supplementary Figure 5**). To compute the value of the KS test statistic, we used the ks_2samp function from the scipy.stats python library.

#### Assessing agreement between HiDDEN continuous perturbation scores and ground truth labels

To quantify the ability of the continuous perturbation scores output by HiDDEN to capture the biological difference between Memory B and Naive B cells in each simulated dataset, we computed the Area Under the Receiver Operating Characteristic Curve (AUROC) using the roc_auc_score function from the sklearn.metrics python library (**Figure 2F, Supplementary Figures 4A, 5B, E**). The AUROC score can take values in the range [0, 1], with higher values indicating better agreement between the continuous prediction scores and the ground truth Naive B / Memory B binary labels.

#### Assessing agreement between HiDDEN refined binary labels and ground truth labels

To evaluate the agreement between ground truth Naive B / Memory B labels and the binary labels refined by HiDDEN in each simulated dataset (**Supplementary Figure 4B**), we used the mutual information score as a typical metric of the similarity of two labels of the same data. To compute the mutual information score, we used the mutual_info_score function from the sklearn.metrics python library. The value of the mutual information score depends on the total number of samples in a dataset. To enable comparison of the mutual information score across all ground truth datasets containing a variable total number of cells, we standardized the value per dataset by the mutual information score computed between the ground truth labels with themselves. The standardized mutual information score can take values in the range [0, 1], with higher values indicating better agreement between the HiDDEN refined binary labels and the ground truth Naive B / Memory B binary labels.

#### Measuring accuracy of retrieved ground truth markers

Here we consider two scenarios – (1) working with the (unclustered) dataset as a whole and (2) looking for perturbation markers across all clusters produced by the standard Seurat workflow.

The unclustered case: To define the set of ground truth markers for Naive B and Memory B cells in each simulated dataset, we computed the DE genes using the Naive B / Memory B ground truth labels using the Wilcoxon rank-sum differential expression test in scanpy with adjusted p-value threshold < 0.05. Analogously, we defined the set of HiDDEN marker genes and a baseline set of marker genes using the refined binary labels and the sample-level case-control labels, respectively, under the same testing procedure. We then computed the number of True Positives (TP), False Negatives (FN), False Positives (FP), and associated metrics of Recall and

Precision (**Figure 2G, Supplementary Figures 2, 3, 4**). Recall can take values in the range [0, 1], with higher values indicating a higher fraction of correctly retrieved ground truth markers. Precision can take on values in the range [0, 1], with lower values indicating a higher fraction of falsely discovered marker genes.

The clustered case: We proceeded analogously to the unclustered case with the difference that DE testing was performed per cluster and the union of all DE genes across clusters defined the final gene set (**Supplementary Figure 2**).

#### Comparing HiDDEN to CNA, MELD, and Milo

While these four methods are developed with different objectives within the larger question of characterizing the effect of a perturbation in single-cell data, all four of them utilize expression profiles (or a neighborhood graph derived from them) and cell labels reflecting the condition of the sample a cell comes from (as well as other metadata, optionally as input. All four methods output a continuous score measuring the effect of the perturbation in each cell, which can further be binarized whenever appropriate. Therefore, we can compare the performance of these four methods along both continuous scores and binary labels. Towards that end, we use the ground truth datasets of Naive B and Memory B cell mixtures.

Training of CNA was performed in python following the jupyter notebook tutorial provided by the authors of the method^32^. The original CNA implementation has a hard-coded assumption that the dataset to be analyzed is composed of at least five samples. We relaxed this assumption to accommodate our B cell mixtures, and obtained continuous perturbation scores as the per-cell neighborhood coefficient using the CNA association function with case/control status as the sample-level attribute of interest and case/control status as sample id. The resulting CNA continuous score is a correlation value ranging from -1 to 1.

Training of MELD was done in python following the code used by the authors of Milo in their comparison section^33^. We computed the k-nearest neighbors graph based on the expression matrix subsetted to the top 2000 highly variable genes. Then we used the meld_op.transform function to compute the density of the case/control sample labels and transformed the density to likelihood per condition using the meld.utils.normalize_densities function. Continuous perturbation scores were obtained as the likelihood of label 1 which takes on values between 0 and 1.

Training of Milo was performed in python following the Differential abundance analysis in python with milopy jupyter notebook tutorial provided by the authors of the method^34^. The underlying implementation of Milo is done in R and calls on the glmFit routine from the edgeR R package. This routine cannot estimate the negative binomial dispersion parameter if it is not given at least three samples. To overcome this hard-coded assumption, we randomly created three samples per condition. We created the partially overlapping cellular neighborhoods using the milo.make_nhoods function. This results in a collection of neighborhoods, each identified by an index cell. A fraction of the cells in the dataset are deliberately excluded from the Milo analysis as outliers. Cells included in the analysis can also belong to one or more neighborhoods at the same time. Then we used the case/control sample level labels to count the number of cells from each sample in each neighborhood using the milo.count_nhoods function. We used the milo.DA_nhoods function to perform differential abundance testing, which outputs a log fold-change test statistic per neighborhood. Since we aim to make comparisons at the level of individual cells, when a cell belonged to more than one neighborhood - we reported the average log fold-change across neighborhoods. For cells excluded from the analysis, the average log fold-change is NA. Continuous perturbation scores were obtained as the average log fold-change and can take on values between minus to plus infinity.

Comparison of the continuous perturbation scores from HiDDEN, CNA, MELD, and Milo (**Supplementary Figure 5A, Supplementary Figure 6**) was performed using the AUROC described earlier in the Methods section as a metric for assessing agreement between continuous perturbation scores and ground truth labels.

Converting the continuous perturbation scores into binarized labels was done in identical manner for all four methods, as described earlier in the Methods section. Comparison of the binary labels from HiDDEN, CNA, MELD, and Milo (**Supplementary Figure 5B, Supplementary Figure 7**) was performed using the Standardized mutual information described earlier in the Methods section as a metric for assessing the agreement between HiDDEN-refined binary labels and ground truth labels.

#### Using HiDDEN-refined binary labels as input to CNA, MELD, and Milo

HiDDEN-refined binary labels demonstrate better agreement with ground truth labels than the sample-level case/control labels. Therefore, we explored the performance of CNA, MELD, and Milo when given HiDDEN binary labels in lieu of case/control labels as input (**Supplementary Figure 6, 14**). CNA continuous perturbation scores were obtained as the per-cell neighborhood coefficient using the CNA association function with Memory B / Naive B ground truth labels as the sample-level attribute of interest and HiDDEN-refined binary labels as sample id. MELD continuous perturbation scores were obtained as the density of the HiDDEN-refined binary labels transformed to likelihood of label 1. Milo continuous perturbation scores were obtained as the average log fold-change downstream of differential abundance testing per neighborhood using the HiDDEN-refined binary labels to count the number of cells per condition. All continuous scores were binarized in the same manner as above.

### Assessing performance on human multiple myeloma and precursor states data

The first real dataset we analyzed consists of single-cell RNA-seq profiles of human bone marrow plasma cells from patients with multiple myeloma (MM) (*n* = 8 patients, *N* = 10790 cells), its precursor conditions smoldering multiple myeloma (SMM) (*n* = 12 patients, *N* = 8431 cells) and monoclonal gammopathy of undetermined significance (MGUS) (*n* = 6 patients, *N* = 817 cells), and healthy donors with normal bone marrow (NBM) (*n* = 9 patients, *N* = 9329 cells)^4^.

Precursor samples can contain a mixture of neoplastic and normal cells (**Figure 3A**) and the authors of the original study define two sources of ground truth describing the malignancy status of precursor samples and their cells. The first source is a manual annotation of binary labels reflecting whether a cell is healthy or malignant. The second source is a tumor-purity estimate of the proportion of malignant cells in each precursor sample.

#### HiDDEN model training

This dataset has pronounced patient-specific batch effects (**Supplementary Figure 8A**). Therefore, echoing the analysis in the original study, we first deployed a batch-sensitive strategy to refine the malignancy status of cells in precursor samples using HiDDEN. Additionally, we developed a batch-agnostic strategy as well, to explore the ability of our method to perform well in the presence of strong batch effects. Mirroring the within-patient annotation approach in the original study, our batch-sensitive fitting approach considers each precursor sample one at a time. We trained the model on all NBM, one precursor sample, and all MM samples, where we give all NBM cells label 0, or healthy, and all of the rest label 1, reflecting that they do not originate from healthy donors. The batch-agnostic fitting approach consists of fitting the model to all NBM samples, all precursor samples manually annotated to be mixed, and all MM samples together. Besides this, all other aspects of model training were carried out the same way between the two strategies. The results from the batch-specific strategy are featured in **Figure 3**, and the results of the batch-agnostic approach, along with a comparison of the two, are included in **Supplementary Figure 8**.

The HiDDEN model training was done in python. First, we followed the standard preprocessing routine for single-cell RNA-seq data in scanpy and log-normalized each sample separately. We then used the standard PCA dimensionality reduction routine in scanpy on the scaled gene features. Next, we used the LogisticRegression function from the sklearn.linear_model python library to train a logistic regression on the binary NBM / non-NBM labels and the first *P* PCs. Finally, we used the KMeans function from the sklearn.cluster python library with *n_clusters* = 2 on the continuous perturbation scores output by the logistic regression for cells in each precursor sample separately (**Supplementary Table 2**). We used all genes to compute the PC dimensionality reduction and automatically chose *P*, the number of PCs used in the logistic regression, using the heuristic for maximizing the number of DE genes downstream of the HiDDEN refined binary labels (**Supplementary Figure 9**). The specific strategy for defining the DE genes in this dataset characterized by strong batch effects is described in detail below.

Under both fitting strategies, we used the same downstream metrics to evaluate the performance of our method at recovering the manually annotated cell-level labels and the purity sample-level estimates, as described below.

#### Assessing agreement between HiDDEN continuous perturbation scores and ground truth manually annotated labels

To quantify the ability of the continuous perturbation scores produced by HiDDEN to capture the manually annotated healthy-malignant binary labels in all mixed precursor samples, we computed the AUROC using the roc_auc_score function from the sklearn.metrics python library. The average AUROC across samples per precursor state from the batch-sensitive training strategy is depicted in **Figure 3B**, and the per-sample curves and distributions of perturbation scores are included in **Supplementary Figure 6**.

#### Assessing agreement between HiDDEN refined binary labels and tumor-purity sample estimates

The sample-level source of ground truth provided in the original study is an estimate of the proportion of malignant cells from a Bayesian hierarchical model based only on the expression of immunoglobulin light chain genes. Additionally, we estimated the per-sample tumor-purity using the manual labels and the HiDDEN refined binary labels. Confidence bounds in all three cases were derived following the same approach as in the original study (**Figure 3C, E**).

#### Differential expression analysis using manually annotated and HiDDEN binary labels

To demonstrate the ability of the HiDDEN-refined binary labels to discover additional malignancy markers from precursor samples, we computed the DE genes using the manual annotation in NBM and MM samples, and contrasted it against the DE genes found using the HiDDEN refined labels in precursor samples (**Figure 3D**). Due to the presence of strong batch effects in the data, mirroring the DE testing strategy in the original paper, we find the DE genes per patient and take the union across patients. For a gene to be considered DE, it had to have an adjusted p-value < 0.05 from the t-test differential expression testing routine in scanpy and a maximum absolute log-foldchange > 1.5.

To assess the significance of the overlap between the two sets of DE genes, we ran a hypergeometric test using the dhyper function from the Stats R package. The background number of genes was calculated based on genes expressed in at least one cell from all precursor samples (18, 770 genes).

#### Validation of HiDDEN binary labels in non-mixed MGUS samples

The authors of the original study computed a Bayesian non-negative matrix factorization (NMF) to highlight gene signatures that are active in this patient cohort and validated them in external cohorts. Several signatures were annotated with a biological interpretation. There were three MGUS samples considered to consist of only healthy cells, according to the manual annotation. They are also the three precursor samples with lowest, although not zero, estimated sample purity according to the Bayesian purity model from the original study. The HiDDEN refined binary labels for these patients annotate some of their cells as malignant. To validate this annotation, we plotted the mean activity of the genes identified by the original study with each signature in the cells labeled as healthy and as malignant in each sample (**Figure 3F, Supplementary Figure 7**). The confidence bounds in **Figure 3F** were derived following the same approach as in the original study.

### Analysis of mouse endothelial cells from a time-course demyelination experiment

#### Generation of demyelination and control tissue

The second real dataset analyzed consists of single-nucleus RNA-seq profiles of mouse endothelial cells from a demyelination model with matched controls. Case and control animals received 500nl of 1% lysophosphatidylcholine (LPC, Cat# 440154, Millipore Sigma, US) or saline vehicle (PBS) injection, respectively. This was delivered using intracranially, using standard approaches, with a Nanoject III (Drummond, US) into the corpus callosum at the following stereotaxic coordinates: Anterior-Posterior: -1.2, Medio-lateral: 0.-5 relative to bregma and a depth of 1.4mm normalized to the surface of the skull.

Mice were sacrificed at four time points: 3, 7, 12, and 18 days post injection (dpi) with *n* = 3 animals per time point per condition, totaling *n* = 24 animals (**Figure 4A**). At the appropriate time point, mice were perfused with ice-cold pH 7.4 HEPES buffer (containing 110 mM NaCl, 10 mM HEPES, 25 mM glucose, 75 mM sucrose, 7.5 mM MgCl2, and 2.5 mM KCl) to remove blood from the brain. Brains were fresh frozen for 3 min in liquid nitrogen vapor and all tissue was stored at −80 °C for long-term storage. The full dataset will be described in a forthcoming paper (Dolan et al, in preparation).

#### Generation of single-nucleus RNA profiles

Frozen mouse brains were mounted onto cryostat chucks with OCT embedding compound within a cryostat. Brains were sectioned until reaching the injection site location, which was confirmed by the presence of hypercellularity using a Nissl stain (Histogene Staining Solution, KIT0415, Thermofisher). For saline controls, anatomical landmarks were used to determine the injection site. Lesions or control white matter was microdissected using a 1mm biopsy punch (Integra Miltex, US), whose circular punch was bent into a rectangle shape with a sterile hemostat. Lesion punches were 300 *μ*m deep.

Each excised tissue punch was placed into a pre-cooled 0.25 ml PCR tube using pre-cooled forceps and stored at −80 °C for a maximum of 24 hours. Nuclei were extracted from this frozen tissue using gentle, detergent-based dissociation, according to a protocol available at protocols.io (https://doi.org/10.17504/protocols.io.bck6iuze) with minor changes to maximize nuclei extraction, which will be described in a forthcoming paper (Dolan et al. in preparation). Nuclei were loaded into the 10x Chromium V3 system. Reverse transcription and library generation were performed according to the manufacturer’s protocol (10x Genomics). Sequencing reads from mouse cerebellum experiments were demultiplexed and aligned to a mouse (mm10) premrna reference using CellRanger v3.0.2 with default settings. Digital gene expression matrices were generated with the CellRanger count function. Initial analysis and generation of overall UMAP and clustering (**Figure 4B**) was performed with Seurat v3^25^.

#### Clustering analysis of endothelial cells

For the UMAP representations in **Figure 4C, D**, and **Supplementary Figure 10** of the profiles of *N* = 891 endothelial cells from *n* = 6 animals spanning both case and control conditions and all four timepoints, we used the standard scanpy functions to log-normalize and scale the gene counts, ran PCA, computed the nearest-neighbor graph with 10 neighbors in the latent space defined by the first 50 PCs, and ran the UMAP algorithm.

To cluster the endothelial cells (**Figure 4D**), we used the standard preprocessing and clustering workflow in Seurat. We normalized the gene counts, performed variable gene selection with default parameter values, scaled the normalized counts, performed dimensionality reduction using PCA with default parameter values, built the nearest-neighbor graph with the default number of nearest neighbors, and used the Leiden algorithm with resolution parameter = 0.5 to find clusters.

For the heatmaps in **Figure 4E** and **Supplementary Figure 10**, we computed the abundance of case (LPC) and control (PBS) labels in each cluster and across time points, respectively.

#### HiDDEN model training

Model training was performed in python on all *n* = 891 endothelial cells together. We followed the standard preprocessing routine for single-cell RNA-seq data in scanpy and log-normalized the gene counts, followed by PCA dimensionality reduction of the scaled gene features. For the continuous perturbation score in **Figure 4F** and **Supplementary Figure 10**, we used the LogisticRegression function from the sklearn.linear_model python library to train a logistic regression on the binary PBS / LPC labels and the first *P* PCs. We used all genes to compute the PC embedding and automatically chose *P* = 5, the number of PCs used in the logistic regression, using the heuristic for maximizing the number of DE genes downstream of the HiDDEN refined binary labels. The strategy for defining the DE genes in this dataset is described in detail below.

For the HiDDEN refined binary labels in **Figure 4G**, we used the KMeans function from the sklearn.cluster python library with *n_clusters* = 2 to split the continuous perturbation scores of all LPC endothelial cells at 3 dpi into two groups. We denote the group with lower perturbation scores LPC0, corresponding to endothelial cells unaffected by the LPC injection and similar to endothelial cells in the PBS control condition, and the group with higher perturbation scores LPC1, as the subset of endothelial cells affected by the LPC injection.

#### Differential expression analysis to define endothelial LPC1 markers

To define the set of endothelial LPC1 markers depicted in the dotplot in **Figure 5A**, we took the unique perturbation-enriched genes found in DE analysis using the HiDDEN refined labels and not in DE analysis using the original PBS / LPC labels. We performed both DE analyses using the Wilcoxon rank-sum test in scanpy with a threshold for the adjusted p-value < 0.05. The comprehensive output from both tests can be found in **Supplementary Table 3**, with the unique genes found using the HiDDEN refined binary labels highlighted in bold font.

#### Validation of endothelial LPC1 markers using RNAscope

Fresh-frozen, 14 *μ*m sections of 3 days post injection (dpi) demyelinating or saline control tissue were mounted on cold Superfrost plus slides (Fisher Scientific, US). These slides were stored at −80 °C. We performed RNAscope Multiplex Fluorescent v2 (Advanced Cell Diagnostics, US) using probes targeting S100a6 (412981), Lgals1 (897151-C2) and Flt1 (415541-C3), where Flt1 is a general marker for endothelial cells. RNAscope was performed following the manufacturer’s protocol for fresh frozen tissue and the following dyes were used at a concentration of 1/1500 to label specific mRNAs (TSA Plus fluorescein, TSA Plus Cyanine 3, TSA Plus Cyanine 5 from PerkinElmer, USA). Imaging was performed on an Andor CSU-X spinning disk confocal system coupled to a Nikon Eclipse Ti microscope equipped with an Andor iKon-M camera. Images were acquired using 20x air and 60x oil immersion objectives (Nikon). All images shown in **Figure 5B** are representative images taken from at least 2 independent experiments.

#### Interpretation of endothelial LPC1 markers using gene ontology analysis

For the identification of gene ontology (GO) categories summarizing the list of unique endothelial LPC1 markers (**Figure 5C, D**), we performed GO enrichment analysis g:Profiler^35^ with default settings and ReviGo^36^ to summarize and visualize the results.

#### Ligand-receptor analysis to identify cell-cell communication changes between endothelial LPC1 and LPC0

To contrast the ligand-receptor communication of the two endothelial LPC subtypes with neighboring cell types in the tissue (**Figure 5E**), we used a modification (Goeva et al, in preparation) to CellphoneDB^37^. We separated the output for each interaction in three bins based on the sign of the test statistic (**Supplementary Figure 11**) and the magnitude of the p-value, reflected in the figure legend: significantly depleted in LPC1 with respect to LPC0, not significant (p-value ≥ 0.05), and significantly enriched in LPC1 with respect to LPC0.

## Data availability statement

The endothelial single-nucleus RNA-seq data will be deposited in Gene Expression Omnibus (GEO).

## Code availability statement

Code and scripts to reproduce analyses presented here are available on Github at https://github.com/tudaga/LabelCorrection.

## Authorship contribution statements

AG developed the algorithm and performed all analyses. MJ-D acquired the demyelination data and performed the imaging validation experiments, with help from JL and EG. RB and RMG assisted with biological interpretation of the analyses. AG and EM conceived the study and wrote the paper, with contributions from all authors.

## Acknowledgements

We thank B. Sanchez-Lengeling, V. Kozareva, Y. Pita-Juarez, J. Webber, A. Lawler, Z. Piran, and members of the Macosko lab for helpful discussions.

This work was supported by a BroadIgnite Philanthropic Grant to AG, Open Philanthropy Project Award of the Life Sciences Research Foundation to MJ-D, and NIMH grant 5U01MH124602 to EZM.

## Supplementary Figures

**Supplementary Figure 1:**
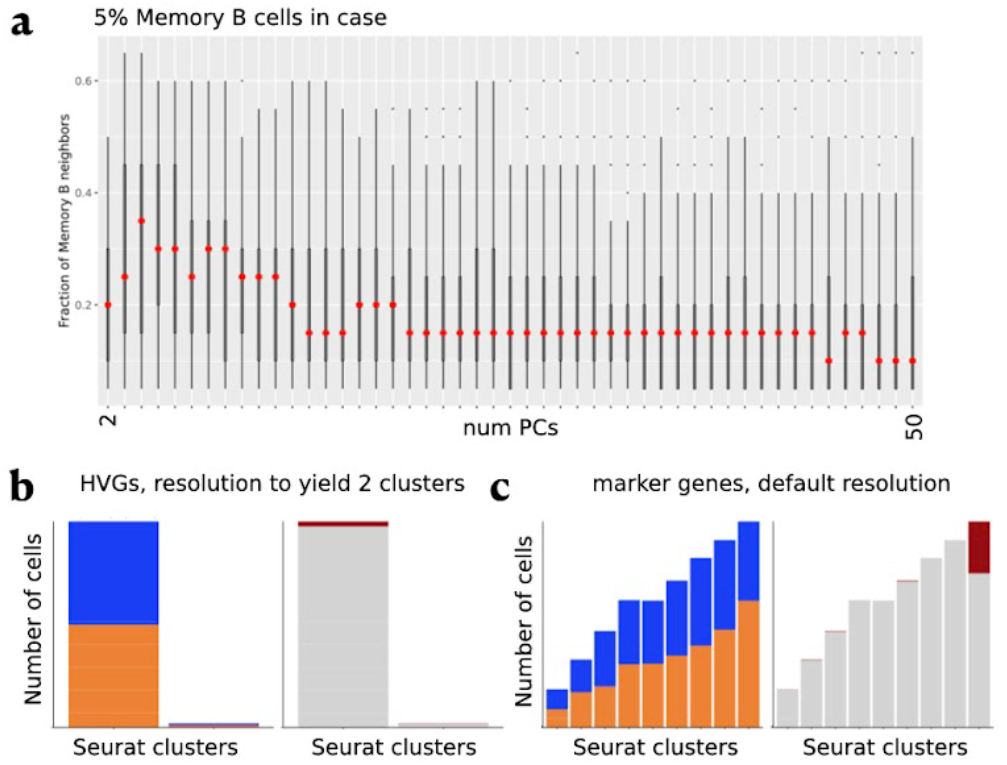
Separability of Naive B and Memory B cells in the latent space and sensitivity of standard dimensionality reduction and clustering workflow to influential parameters. Box plots of the distribution of fraction of Memory B cells among the 20-nearest neighbors of each Memory B cell (y-axis). **A** For the dataset with 5% Memory B cells in case, the fraction of Memory B neighbors in the latent space defined by a variable number of top PCS from 2 to 50 (x-axis). Distribution of case-control (left) and Memory B-Naive B (right) cell identities across Seurat clusters. Colors defined as follows: case-orange, control-blue, Memory B-red, Naive B-gray. Dimensionality reduction and resolution parameters chosen as follows: **B** Using highly variable genes for feature selection (default); resolution parameter chosen to yield two clusters. **C** Using Naive B and Memory B markers for feature selection; default resolution.

**Supplementary Figure 2:**
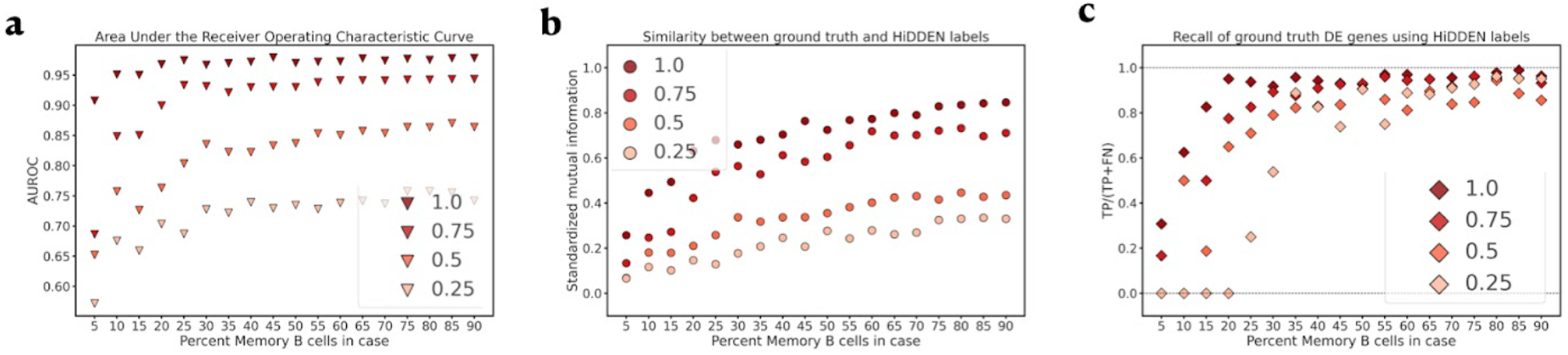
Effect of perturbation strength (as indicated by legend labels) and fraction perturbed cells in case (x-axis) on problem difficulty. y-axis defined as follows: **A** Area under the Receiver Operating Characteristic Curve (AUROC) for classification of ground truth cell labels; **B** Standardized mutual information for assessing agreement between HiDDEN refined binary labels and ground truth labels (Methods); **C** Recall of ground truth DE genes from DE testing on HiDDEN-refined labels.

**Supplementary Figure 3:**
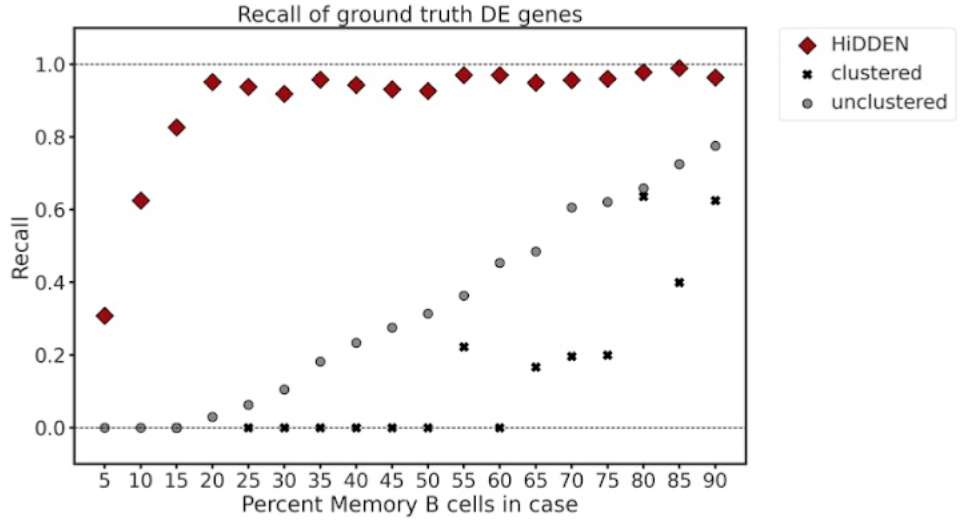
Recall of ground truth DE genes as a function of percent perturbed cells in case sample. Legend labels define the DE testing approach as follows: HiDDEN - using the HiDDEN-refined binary labels on the unclustered data; unclustered - using the case-control labels on the unclustered data; clustered - using the case-control labels to perform DE testing per Seurat cluster and taking the union of DE genes across clusters.

**Supplementary Figure 4:**
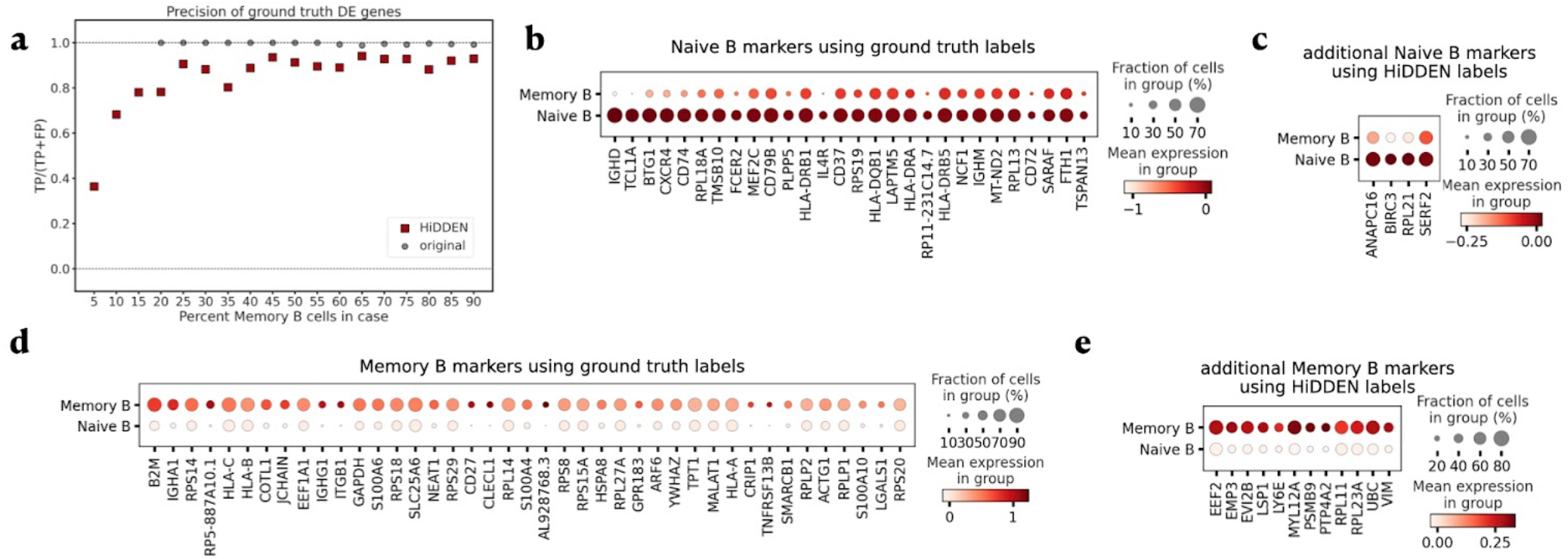
DE genes found from DE testing using HiDDEN-refined binary but not found using Memory B-Naive B labels. **A** Precision of ground truth DE genes from DE testing on HiDDEN-refined and original case-control labels as a function of percent perturbed cells in case sample. Dotplot of mean expression of Naive B (**B**) and Memory B (**D**) marker genes ordered by p-value. Dotplot of mean expression of HiDDEN-refined label 0 (**C**) and label 1 (**E**) DE genes ordered alphabetically.

**Supplementary Figure 5:**
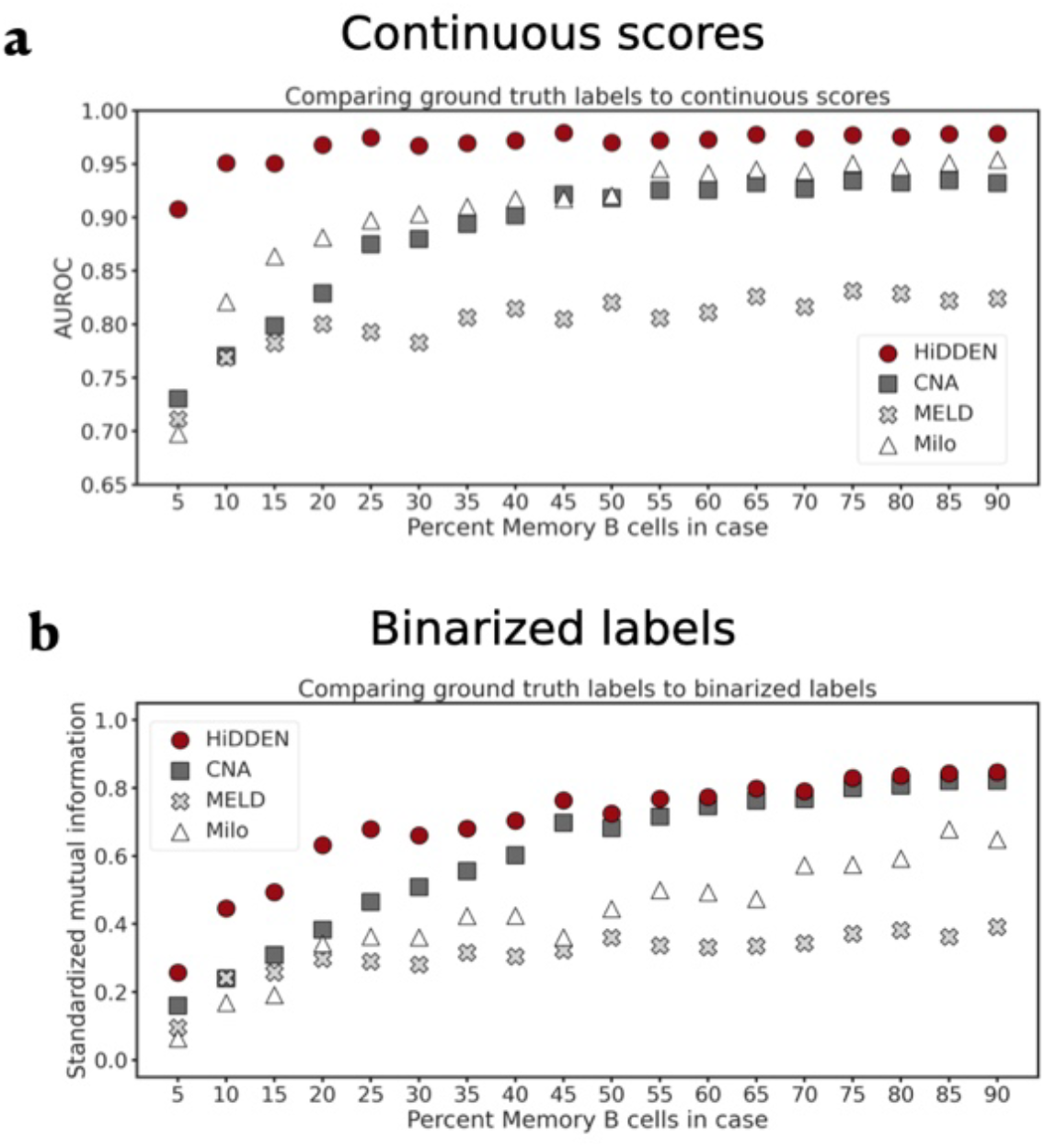
A direct comparison between HiDDEN and CNA, MELD, and Milo. **A** AUROC score for using continuous perturbation scores for classification of ground truth cell labels as a function of perturbed cells in case sample for each of the four methods. **B** Standardized mutual information between ground truth Naive B / Memory B labels and binarized labels as a function of percent perturbed cells in case sample for each of the four methods.

**Supplementary Figure 6:**
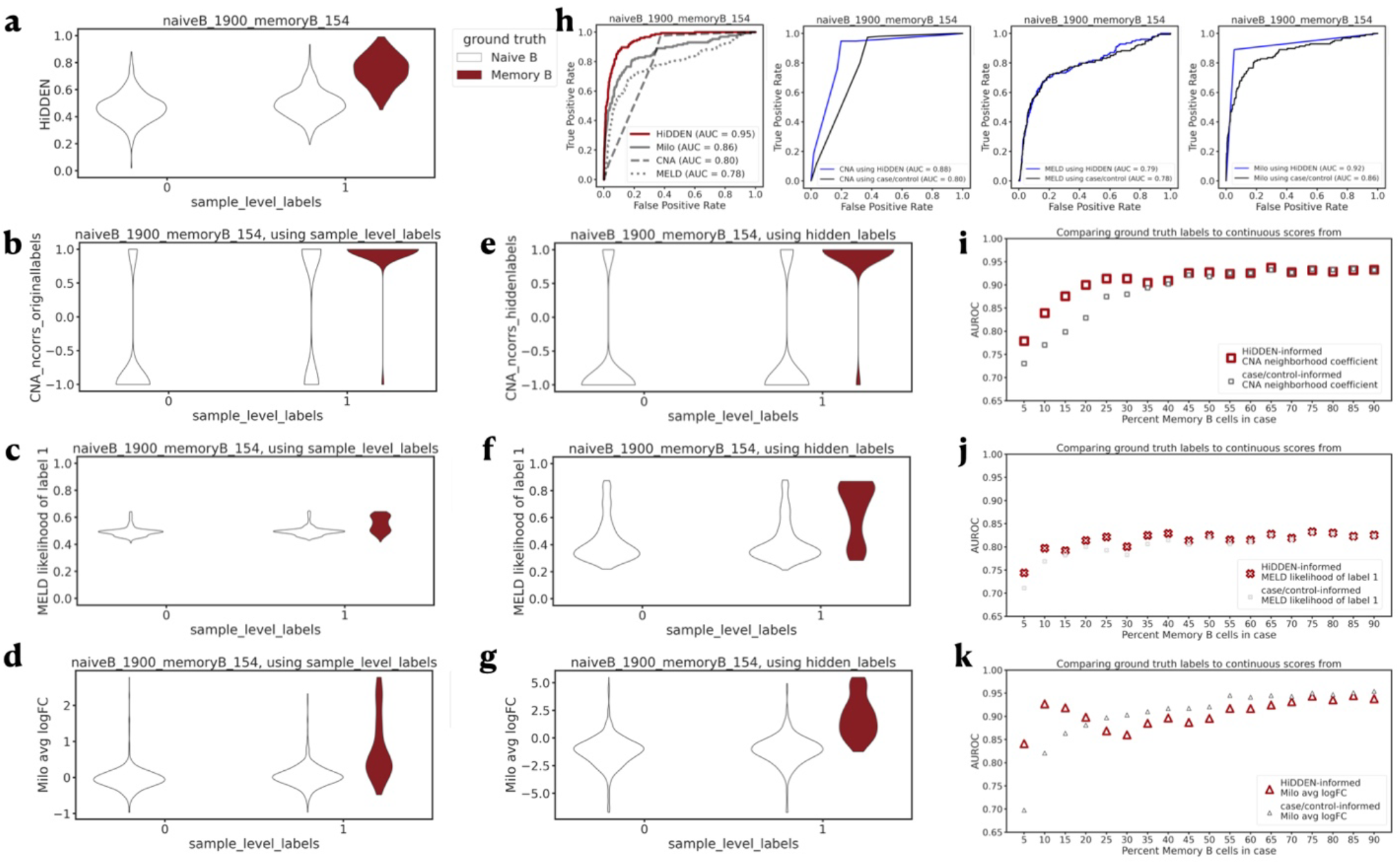
A comparison of HiDDEN-informed vs. case/control-informed performance of CNA, MELD, and Milo continuous scores. Violin plots (area not scaled to count) of the distribution of continuous scores (method and continuous score details detailed on y-axis) of Naive B and Memory B cells split over control and case samples and colored by ground truth labels, for the dataset containing 15% Memory B cells in the case sample. Results in **A, B, C, D** use case/control sample-level labels as input. Results in **E, F, G** use HiDDEN-refined binary labels. **H** Area under the Receiver Operating Characteristic (ROC) curves for classification of ground truth cell labels as a function of percent perturbed cells in case sample with the Area Under the ROC (AUROC) and method indicated in the legend, for the dataset containing 15% perturbed cells in the case sample. **I, J, K** AUROC score for using continuous perturbation scores for classification of ground truth cell labels as a function of perturbed cells in case sample, with the method name reflected in the plot title.

**Supplementary Figure 7:**
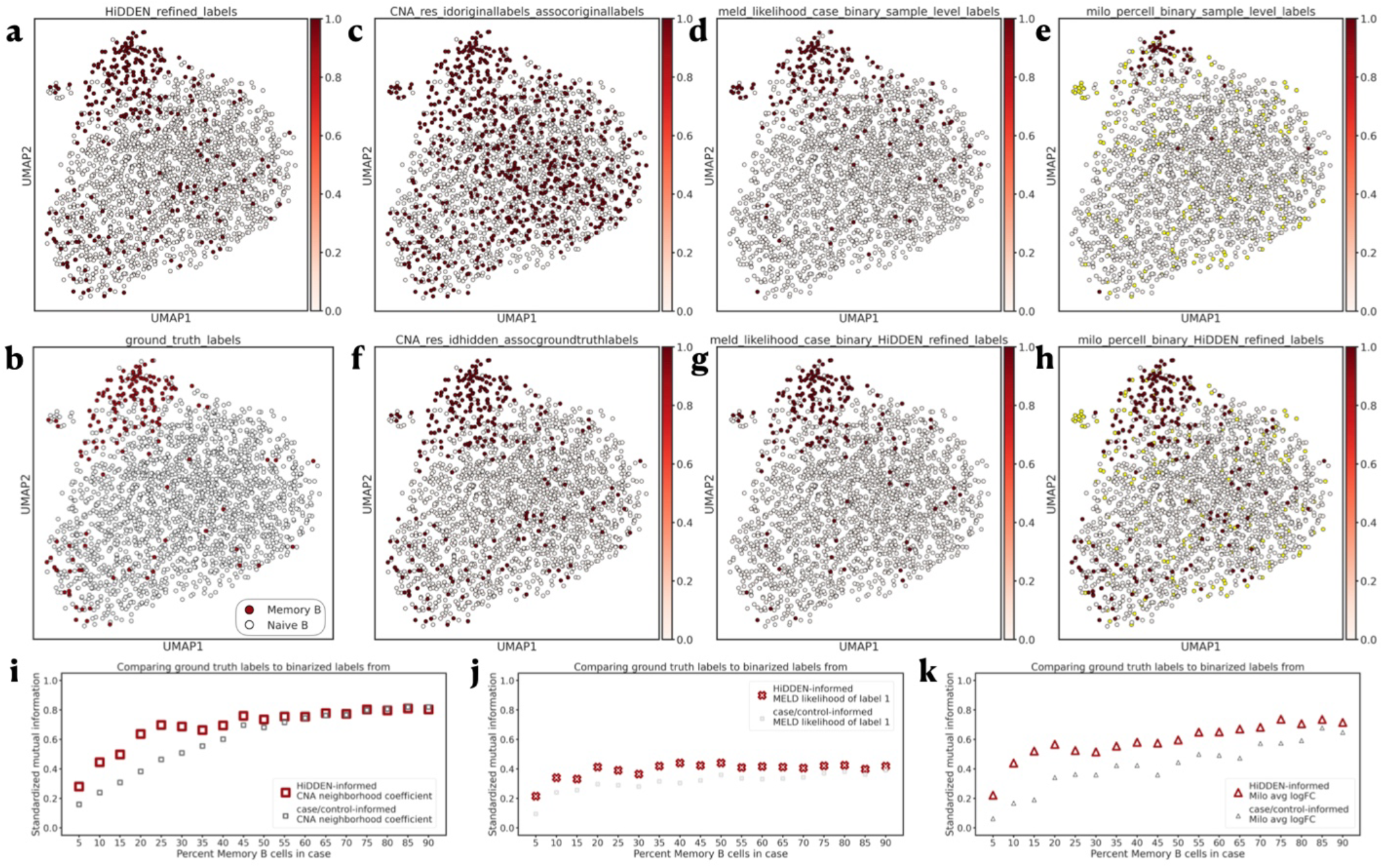
A comparison of HiDDEN-informed vs. case/control-informed performance of CNA, MELD, and Milo binarized labels. UMAP embeddings of Naive B and Memory B cell gene expression for the dataset containing 15% Memory B cells in the case sample colored by: **A** HiDDEN-refined binary labels, **B** ground truth Memory B / Naive B labels, **C, F** CNA binarized neighborhood coefficient, **D, G** MELD binarized likelihood of label Memory B, **E, H** Milo binarized average log fold-change (yellow dots mark NA log fold-change values corresponding to cells excluded by Milo from the analysis); (using case/control or HiDDEN-refined labels, respectively) **I, J, K** Standardized mutual information between ground truth Naive B / Memory B labels and binarized labels as a function of percent perturbed cells in case sample, with the method name reflected in the plot title.

**Supplementary Figure 8:**
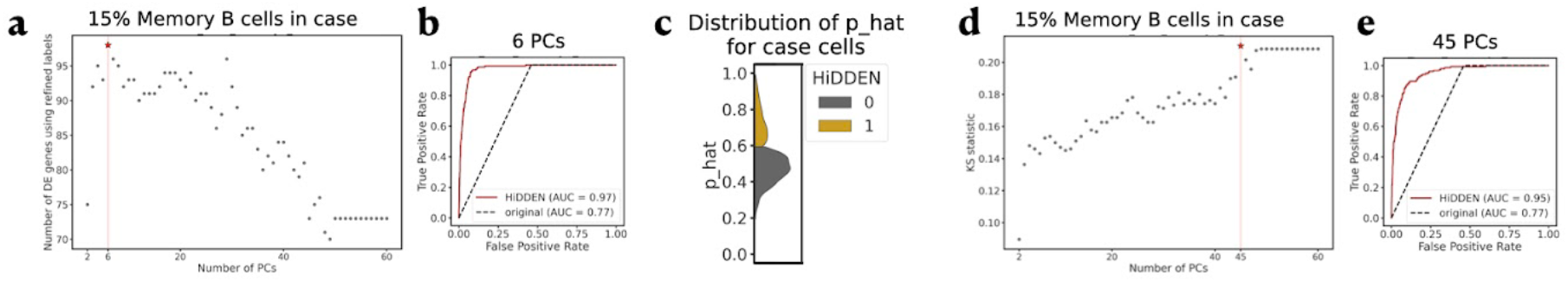
Two heuristics for choosing the number of features (top PCs) for the prediction model part of HiDDEN. For the dataset with 15% Memory B cells in case: **A** Number of DE genes using HiDDEN-refined binary labels (y-axis) as a function of number of PCs (x-axis) with the optimal (6 PCs) marked by a vertical dashed red line and a star. **B** Receiver Operating Characteristic Curve (ROC) for classification of ground truth cell labels of the model using 6 PCs with AUROC and line style and color as defined in the legend. The second heuristic computes the two-sample Kolmogorov-Smirnov statistic (Methods) between the distribution of HiDDEN continuous perturbation scores (p_hat) for HiDDEN-refined binary label 0 and label 1. **C** Density plots of the two distributions with colors as indicated in legend. **D** KS statistic as defined in C (y-axis) as a function of number of PCs (x-axis) with the optimal (45 PCs) marked by a vertical dashed red line and a star. **E** Receiver Operating Characteristic Curve (ROC) for classification of ground truth cell labels of the model using 45 PCs with AUROC and line style and color as defined in the legend.

**Supplementary Figure 9:**
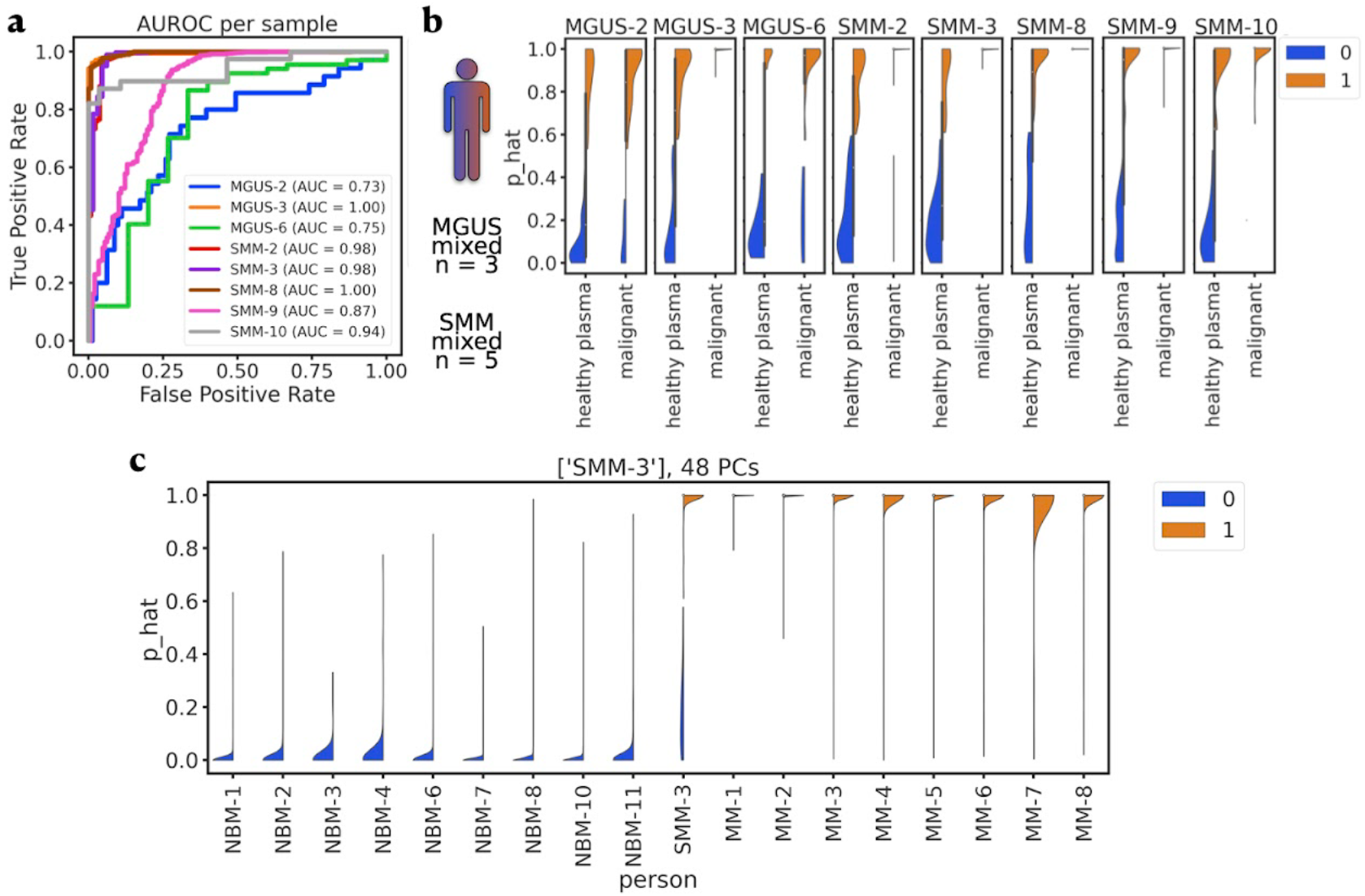
HiDDEN predictions across mixed precursor samples and positive (MM) and negative (NBM) controls. **A** Area under the Receiver Operating Characteristic Curves (AUROC) for each mixed precursor sample plotted individually. **B** Violin plot of the distribution of the continuous perturbation score (y-axis) across cells manually annotated as healthy plasma or malignant (x-axis) split over and colored by HiDDEN-refined binary labels 0 and 1; one panel per mixed precursor sample; area scaled by count. **C** Density plots of the distribution of the continuous perturbation score (y-axis) from the batch-sensitive fitting strategy for the mixed precursor SMM-3 across all samples included in the model (x-axis). Colors correspond to HiDDEN-refined labels; area scaled by count.

**Supplementary Figure 10:**
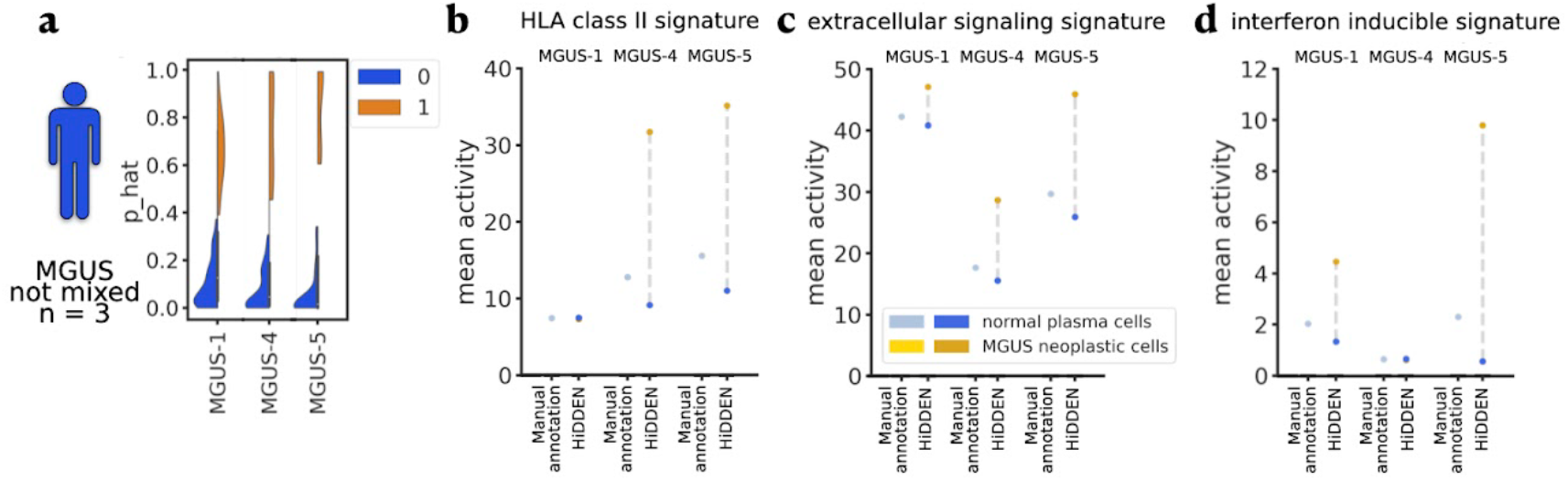
Predictions and computational validation for the low tumor purity MGUS samples considered non-mixed according to the manual annotation. **A** Violin plots of the distribution of the continuous perturbation score (y-axis) across the three low tumor purity MGUS precursors (x-axis) split over and colored by HiDDEN-refined binary labels 0 and 1; area scaled by count. **B, C, D** Computational validation of cells predicted to be malignant by HiDDEN in low tumor purity MGUS samples. Mean activity (y-axis) of genes assigned to three biologically-interpretable signatures from the original study separately computed per HiDDEN label across the three low tumor purity MGUS samples (x-axis).

**Supplementary Figure 11:**
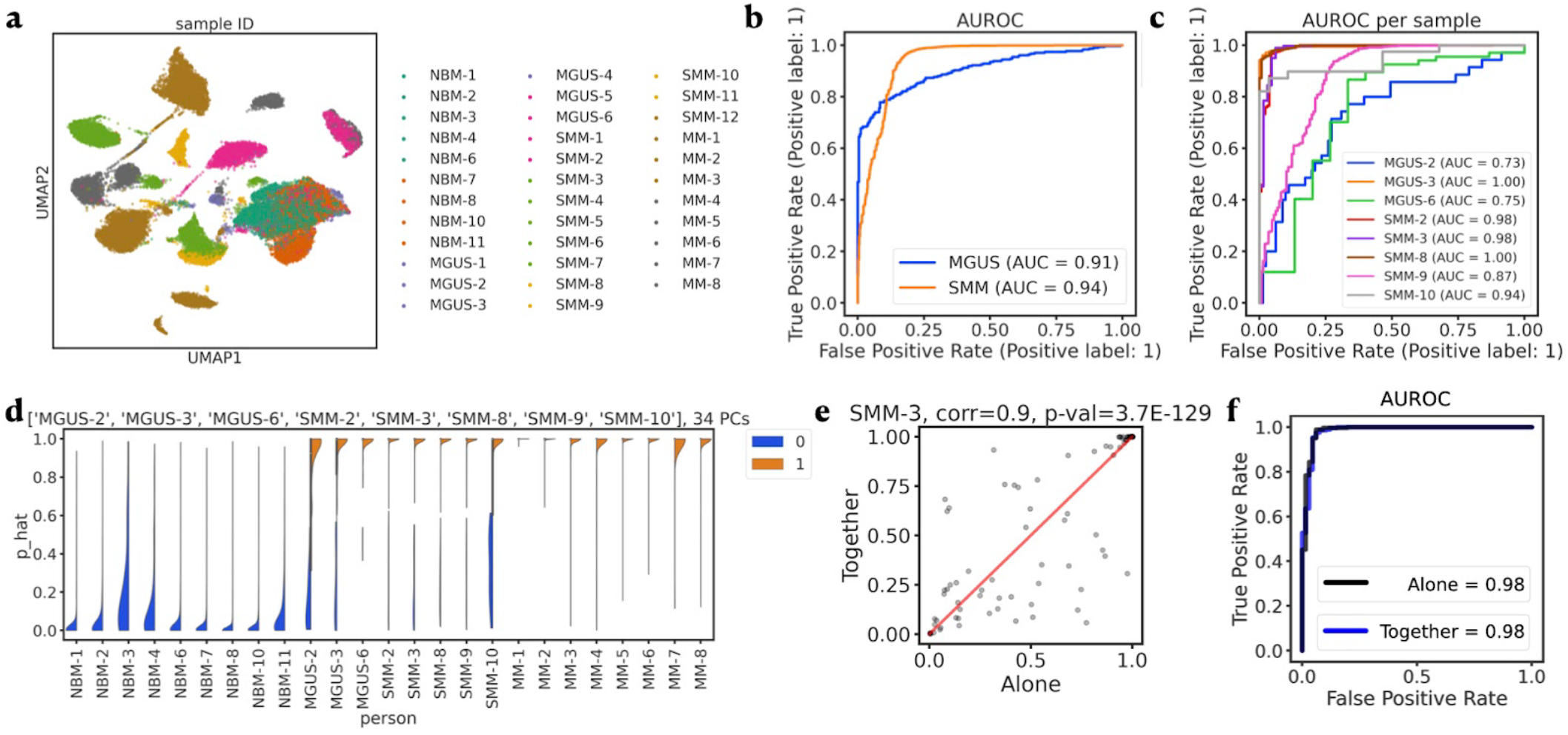
Comparison of batch-sensitive and batch-agnostic fitting strategies for mixed precursor sample SMM-3. **A** UMAP embeddings of human bone marrow cells from all NBM, MGUS, SMM and MM patients from ^4^, colored by patient. **B** AUROC for predicting the per-cell manually annotated malignancy status in mixed samples averaged for each precursor state using the batch-agnostic strategy. Note: Figure 3B shows the results using the batch-sensitive strategy. **C** AUROC for each mixed precursor sample plotted individually using the batch-agnostic strategy. Note: Supplementary Figure 6A shows the results using the batch-sensitive strategy. **D** Density plots of the distribution of the continuous perturbation score (y-axis) from the batch-agnostic fitting strategy for all mixed precursors along with all NBM and all MM across (x-axis). Colors correspond to HiDDEN-refined labels; area scaled by count. Note: Supplementary Figure 6C shows the results using the batch-sensitive strategy for SMM-3. **E** Scatterplot of the per-cell continuous perturbation scores from the batch-sensitive (x-axis) and batch-agnostic (y-axis) strategies for SMM-3 with 45 degree red line, corr=0.9 (p-val=3.7e-129). **F** Comparison of AUROC for SMM-3 with colors corresponding to the two fitting strategies: Alone - batch-sensitive and Together - batch-agnostic.

**Supplementary Figure 12:**
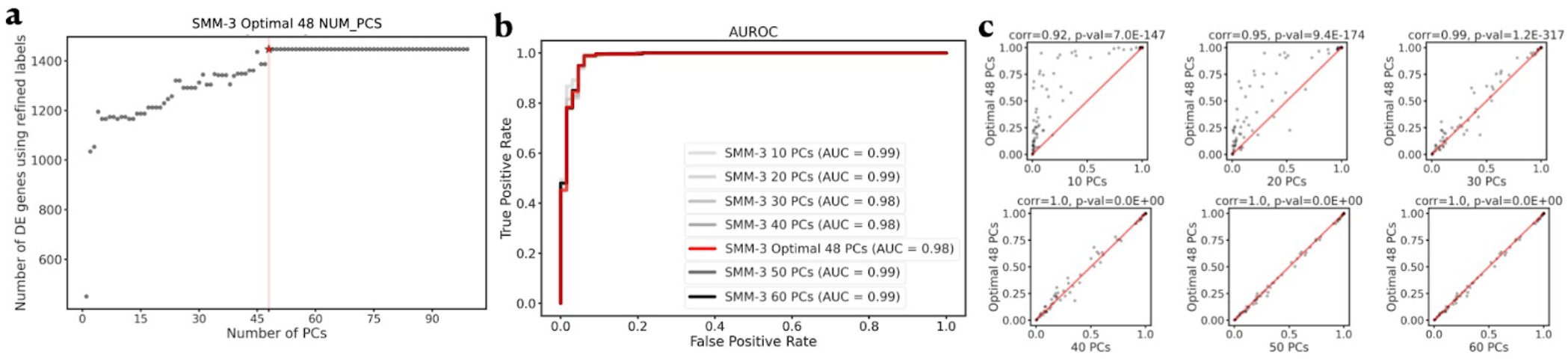
Heuristic for choosing the optimum and sensitivity of the number of features (top PCs) for the bone marrow data illustrated under the batch-sensitive strategy for SMM-3. **A** Number of DE genes using HiDDEN-refined binary labels (y-axis) as a function of number of PCs (x-axis) with the optimal (48 PCs) marked by a vertical dashed red line and a star. **B** ROC curves for classification of manually annotated malignancy cell labels in SMM-3 using a different number of PCs (as indicated by the legend labels). **C** A collection of scatterplots of the per-cell continuous perturbation scores using the optimal number of 48 PCs (y-axis) and each of the number of PCs explored in B (x-axis) with 45 degree red line; correlation and corresponding p-value indicated in the title of each plot.

**Supplementary Figure 13:**
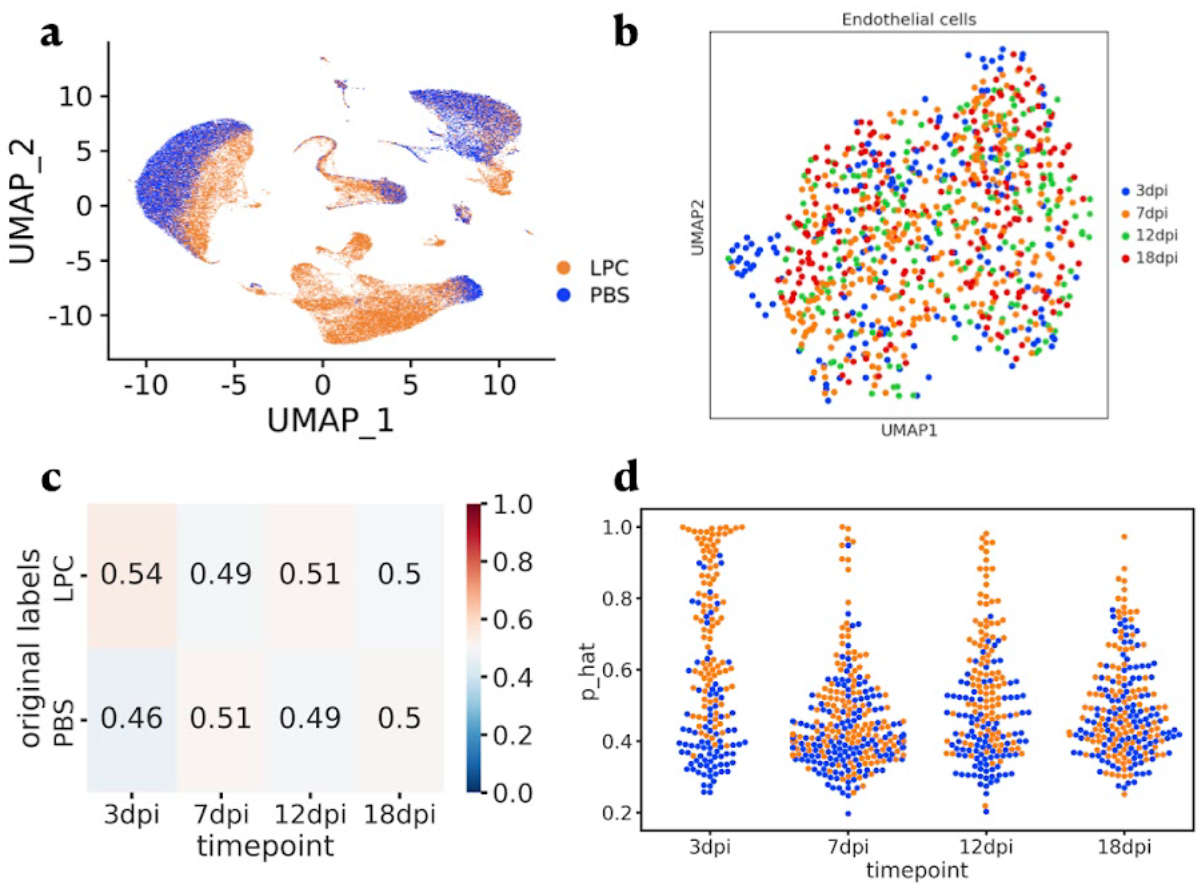
Sample-level and HiDDEN predictions across time points in the mouse demyelination data. **A** UMAP embeddings of non-neuronal cells from both conditions across all time points colored by PBS/LPC sample-level labels. **B** UMAP embeddings of endothelial cells across both conditions colored by time point. **C** Relative abundance of case-control cell identities across time points. **D** Swarmplots of HiDDEN continuous perturbation scores (y-axis) across time points (x-axis); colors as defined in A.

**Supplementary Figure 14:**
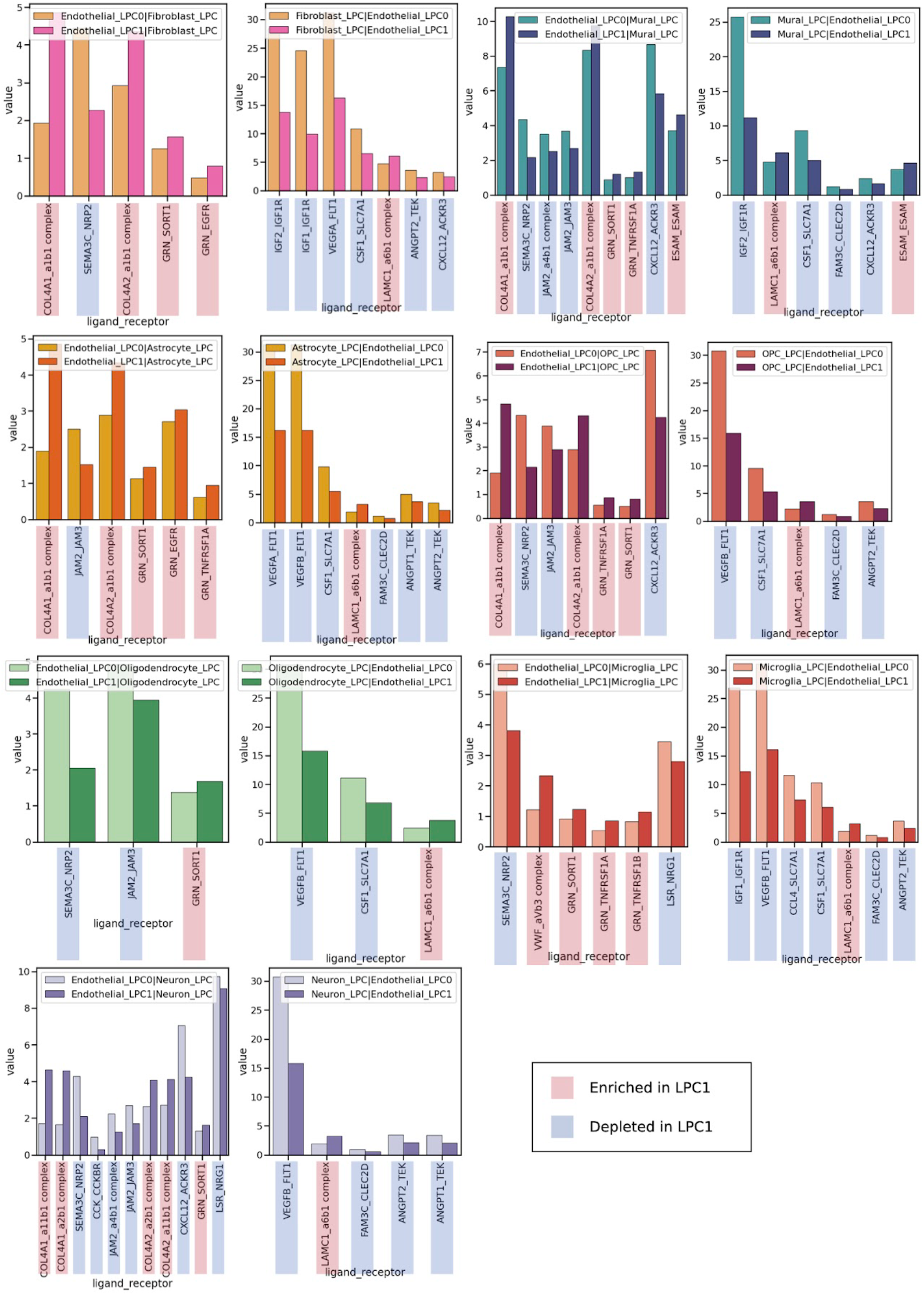
Test statistic for contracting endothelial LPC0 and LPC1 ligand-receptor interactions. A collection of panels each containing a set of plots with two bars. The bars are colored as indicated in each panel’s legend and represent an interaction either from or to one of the endothelial LPC subset and a neighboring cell type. X-axis enlisting the names of significant ligand-receptor pairs, each pair colored to indicate whether it is enriched or depleted. Y-axis is the average of the mean ligand expression and mean receptor expression in their respective cell types.

## Supplementary Tables

**Supplementary Table 1:**
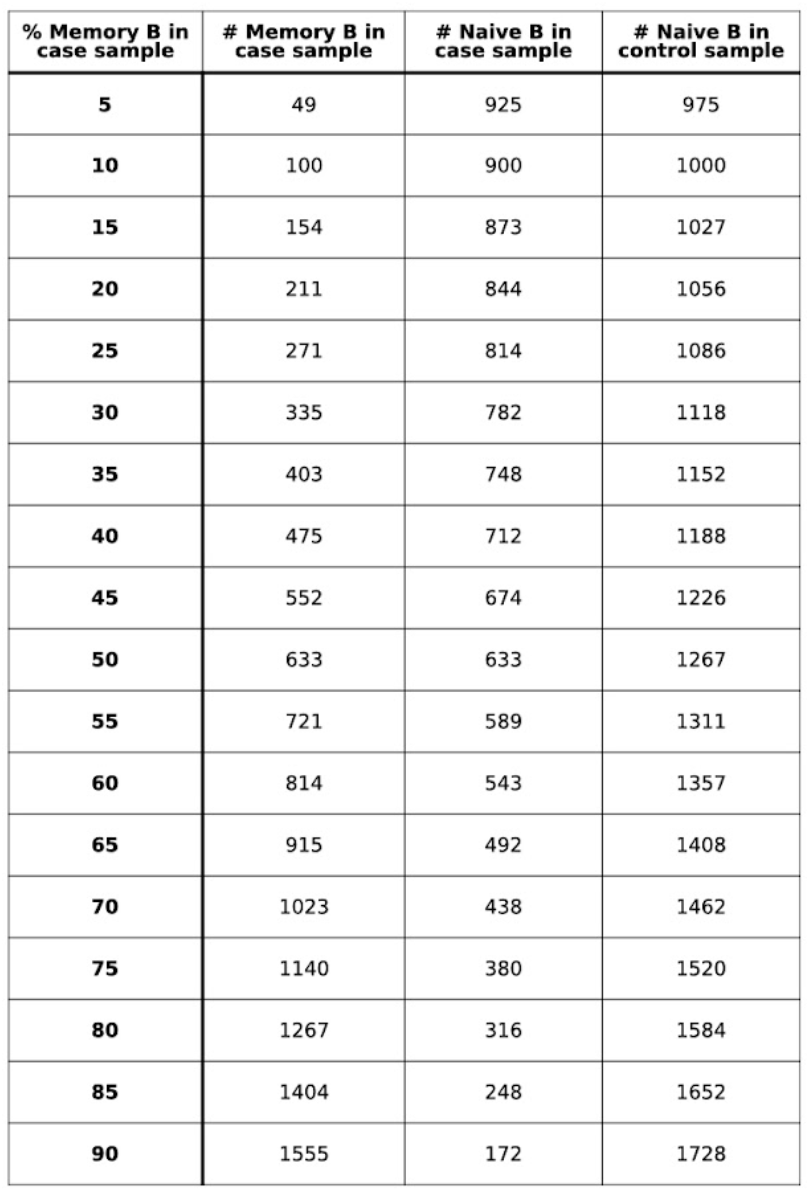
Number of Naive B and Memory B cells in synthetic datasets varying the percent perturbed cells in the case sample. Rows indicate the percent of Memory B cells in case.Columns are: number of Memory B cells in case; number of Naive B cells in case; number of Naive B cells in control.

| % Memory B in case sample | # Memory B in case sample | # Naive B in case sample | # Naive B in control sample |
| --- | --- | --- | --- |
| 5 | 49 | 925 | 975 |
| 10 | 100 | 900 | 1000 |
| 15 | 154 | 873 | 1027 |
| 20 | 211 | 844 | 1056 |
| 25 | 271 | 814 | 1086 |
| 30 | 335 | 782 | 1118 |
| 35 | 403 | 748 | 1152 |
| 40 | 475 | 712 | 1188 |
| 45 | 552 | 674 | 1226 |
| 50 | 633 | 633 | 1267 |
| 55 | 721 | 589 | 1311 |
| 60 | 814 | 543 | 1357 |
| 65 | 915 | 492 | 1408 |
| 70 | 1023 | 438 | 1462 |
| 75 | 1140 | 380 | 1520 |
| 80 | 1267 | 316 | 1584 |
| 85 | 1404 | 248 | 1652 |
| 90 | 1555 | 172 | 1728 |

**Supplementary Table 2:**
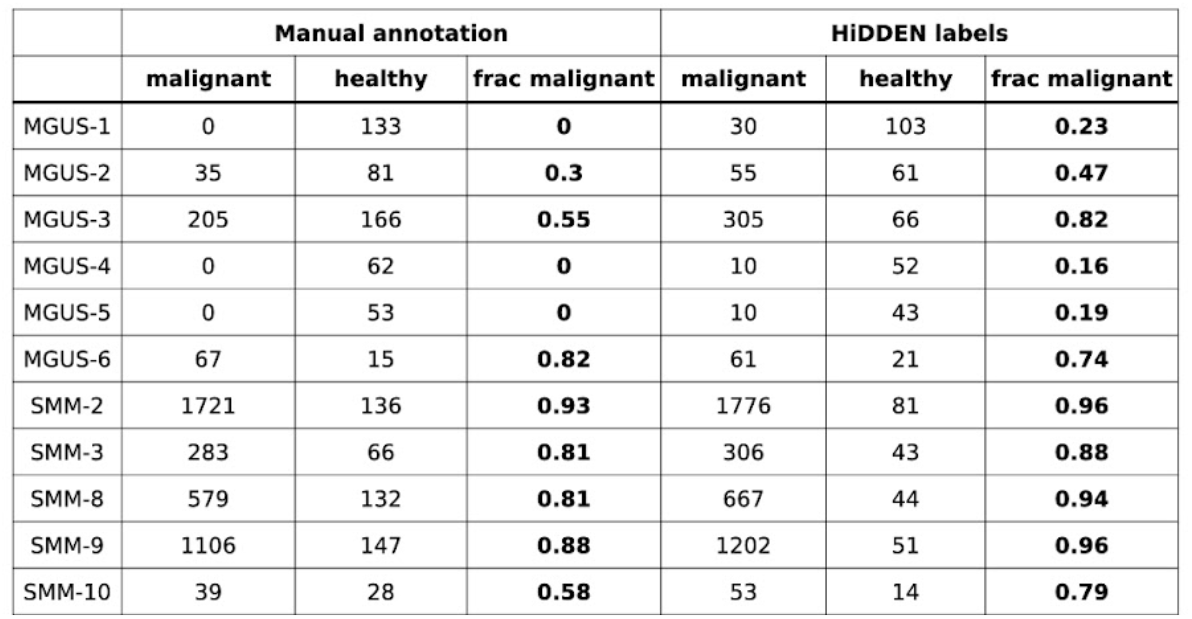
Comparison of manual annotation and HiDDEN outputs across mixed precursor samples. Rows indicate precursor samples labeled as mixed according to the manual annotation. There are two meta-columns: the first reports statistics using the manual annotation, the second reports statistics using the HiDDEN-refined binary labels. Under each meta-column, there are three columns reporting the number of cells annotated as malignant; number of cells annotated as healthy; and the fraction of cells annotated as malignant (to facilitate cross-sample comparison).

**Supplementary Table 3:**
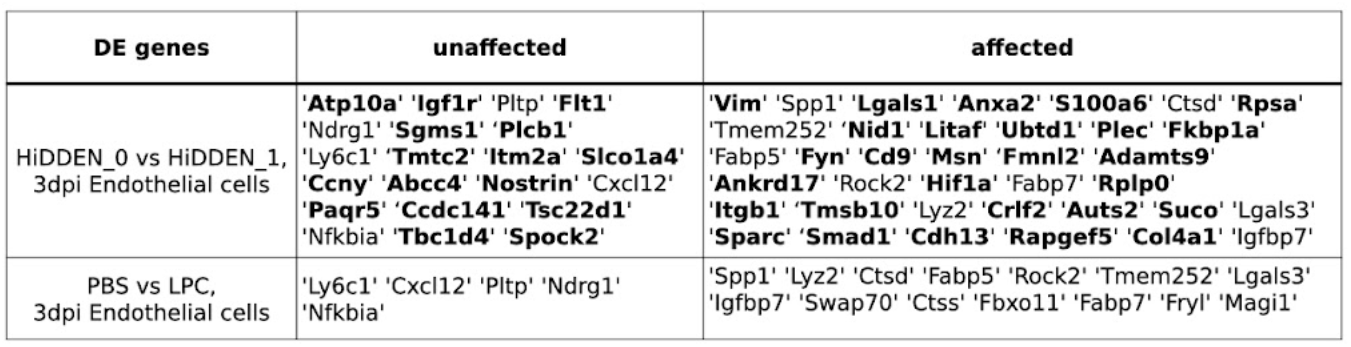
Comparison of DE genes for 3dpi Endothelial cells found using sample-level and HiDDEN labels. Rows indicate labels used in the DE testing (Methods). Columns indicate DE genes for unaffected (associated with HiDDEN_0 or PBS label) and affected cells (associated with HiDDEN_1 or LPC label). Gene names in **bold** are uniquely found using the HiDDEN-refined labels.

